# Genetic Architecture of Addiction-Relevant Behaviors in Outbred Sprague-Dawley Rats Reveals Loci for Anxiety-Like and Nociceptive Traits

**DOI:** 10.64898/2026.02.18.706481

**Authors:** Apurva S. Chitre, Elaine K. Hebda-Bauer, Michael A. Emery, Fei Li, Khai-Minh Nguyen, Yizhi Wang, Riyan Cheng, Oksana Polesskaya, Stanley J. Watson, Jun Li, Huda Akil, Abraham A. Palmer

**Affiliations:** Department of Psychiatry, University of California San Diego, La Jolla, CA, United States; Michigan Neuroscience Institute, University of Michigan, Ann Arbor, MI, United States; Department of Human Genetics, University of Michigan, Ann Arbor, MI, United States; Department of Molecular Genetics and Genome Sciences, University of Oklahoma Health Science Center, College of Medicine, Oklahoma City, OK, United States; Institute for Genomic Medicine, University of California San Diego, La Jolla, CA, United States

## Abstract

Studies have shown that substance use liability is associated with novelty seeking, anxiety-like behavior, and pain sensitivity. We examined whether common genetic variation in outbred Sprague-Dawley rats explained variation in behavioral measures from three assays with established links to substance use: locomotor response to a novel environment, elevated plus maze, and tail flick. We estimated single-nucleotide polymorphism heritability and performed genome-wide association analyses using permutation-derived significance thresholds (N=534-654 rats across traits). Heritability estimates ranged from 0.14-0.38 across eleven traits. Three independent loci were identified: chromosome 1 for elevated plus maze open-arm behavior (α=0.05), chromosome 14 for elevated plus maze immobility (α=0.10), and chromosome 17 for tail flick latency (α=0.05). Candidate genes included *Slc18a2, Gfra1*, and *Pdzd8* (chromosome 1); *Rel* and *Bcl11a* (chromosome 14); and *Eci2* and *Eci3* (chromosome 17). We compared these loci with our genome wide association study of a F_2_ intercross of selectively bred high- and low-responder rats, originally derived from Sprague-Dawleys, that model individual differences in externalizing and internalizing behavior. The current loci are distinct from the ones identified in the bred lines. This difference likely reflects selection history in the high- and low-responder F_2_s, which focused on facets of exploratory locomotion, while loci for anxiety and pain sensitivity traits were identified in the outbreds. This highlights the benefit of using both outbred and selectively bred rats to probe causal variants contributing to individual differences in substance use liability. The current outbred findings implicate monoaminergic signaling, transcriptional control, and lipid metabolism as testable mechanisms for addiction-relevant behaviors.

## 1 Introduction

Substance use disorders place a very large economic and population-health burden. For example, the U.S. economic cost of opioid use disorder plus fatal opioid overdose was estimated at $1.02 trillion in 2017 (Florence et al., 2021). These outcomes co-vary with dimensional liabilities that cut across diagnoses, including both negative emotionality and anxiety and high reactivity to novelty and sensation seeking. Human genetic work shows substantial shared heritable liability between externalizing traits and substance use and related disorders (Karlsson Linnér et al., 2021) and links impulsivity facets to drug experimentation and use (Sanchez-Roige et al., 2019). On the other end of the temperament spectrum, neuroticism, an internalizing trait, is also associated with several loci identified via genome-wide association studies (GWAS) and strong positive genetic correlations with major depressive and anxiety disorders which share co-morbidities with substance use disorders (Sanchez-Roige et al., 2018). Contemporary frameworks further organize these domains and validate them psychometrically in clinical samples (Gunawan et al., 2024). These combined findings suggest that there are multiple paths to substance use disorders, arising from both externalizing phenotypes that can promote drug experimentation, and internalizing phenotypes that can promote the use of drugs to cope with stress, anxiety, and mood dysregulation.

To interrogate these cross-cutting behavioral liabilities mechanistically and genetically, we turned to rats as model organisms. Rats support rich ethology, pharmacology, and causal circuit manipulations, and outbred populations provide levels of linkage disequilibrium (LD) that are amenable to GWAS (Parker and Palmer 2011). Sprague-Dawley rats are widely used in behavioral neuroscience; although they are not extensively used for genetic studies, they have been genetically characterized. Sprague-Dawley rats obtained from Charles River have shown diversity and LD features favorable for GWAS of complex behaviors (Gileta et al., 2022).

We previously performed a GWAS on a F_2_ intercross of our selectively bred high responder/low responder (bHR/bLR) rat lines, which were originally created using Sprague-Dawley founders (Chitre et al, 2023). The bHR/bLR lines differ in their high or low propensity to explore a novel environment and provide a good model for individual differences in temperament and the extremes in human behavioral liabilities. As they diverged with selective breeding for exploratory locomotion, the bHR/bLR lines came to represent the polar extremes of emotionality exhibited by the outbred Sprague-Dawley rats from which they are derived, thus modeling two distinct paths to substance use (Aydin et al, 2021; Clinton et al, 2014; Flagel et al, 2014; Flagel et al, 2016; Stead et al, 2006; Turner et al, 2017). However, given the specific focus of this selective breeding on exploratory locomotion in a novel environment, it is not expected that the bHR/bLR model would capture the whole range of factors that contribute to substance use liability. For example, sensitivity to pain plays a special role in opioid use disorders, and anxiety and negative affect may be especially relevant to the propensity for continued drug use and relapse, sometimes referred to as the “dark side of addiction” (Koob, 2013). Thus, distinct genetic, neural, and behavioral features are likely to play a role in these differing components of substance use disorders and are likely to be differentially detectable based on the population being studied.

To approach this question broadly, we tested whether common genetic variation in commercially outbred Charles River Sprague-Dawley rats contributed to individual differences in response to novelty (novelty-induced locomotion), approach–avoidance anxiety (elevated plus maze, EPM), and noxious thermal nociception (tail flick test), as each of these traits are associated with addiction-relevant behaviors. Specifically, novelty-induced locomotion and anxiety-like behavior are predictors of drug self-administration and relapse-like responding (Flagel et al, 2014; Piazza et al., 1989; Gancarz et al., 2023), and thermal nociception and pain states modulate opioid seeking and consumption (Hipólito et al., 2015; Koob, 2021; Higginbotham et al., 2022). The temperament trait Harm Avoidance in humans (Cloninger, 1986, 1994; Cloninger et al, 1993), which constitutes a higher tendency to feel fear and anxiety accompanied by an elevated emotional response to aversive stimuli, is linked to decreased mu-opioid binding in brain. This reflects not only a biological association between temperament and pain sensitivity but also corresponds with decreased mu-opioid receptor levels in our high-anxiety bLR rats (Turner et al., 2019). Recently, several genes found linked to either basal pain sensitivity or risk for chronic pain conditions (Kringel and Lötsch, 2025) are also widely known to influence the development, maintenance, or severity of addiction (Agarwal et al., 2022; Elman and Borsook, 2016;Tian et al. 2018; Yan et al. 2012). Response latency to noxious thermal stimulation has also been associated with novelty-induced exploratory locomotion, as outbred rats characterized for high and low reactivity to novelty, which are behaviorally selected from the tail ends of the normal distribution of the Sprague-Dawley population, display different basal responses in the tail flick test (Cecchi et al. 2008).

Our goals were to identify allelic associations with these traits across independent Sprague-Dawley-derived populations and to discover additional variants not captured by selective breeding. To this end, we performed GWAS in outbred Sprague-Dawley rats and situated these analyses alongside prior mapping in an F_2_ intercross of bHR/bLR rats in the same broader background, which identified loci for novelty-induced exploratory locomotion and anxiety-like traits (Chitre et al., 2023; Hebda-Bauer et al, 2025). Our findings shed light on genetic differences associated with comparable exploratory locomotion and anxiety-like behaviors in outbred Sprague-Dawley rats vs. selectively bred rats on the same background.

## 2 Methods

### 2.1 Phenotyping

Sprague-Dawley outbred rats were purchased from Charles River Laboratories, Raleigh, North Carolina, USA. Non-cross-fostered, non-littermates arrived at the Michigan Neuroscience Institute, University of Michigan, at 10 weeks of age, approximately 70 days old. They were housed two per cage, with males and females housed in separate rooms. **Supplementary Table S1** includes demographic information about the 654 rats used in this study. Animals were provided ad lib water and standard rat chow (5LOD, PicoLab Laboratory Rodent Diet, LabDiet) in a temperature (70–78^°^F) and humidity (30%– 50%)-controlled room with a 12/12 light/dark cycle (lights on at 06:00, lights off at 18:00). Behavioral testing took place between 07:30 – 13:00 (Zeitgeber Time (ZT) ZT1.5 to ZT7). All procedures were conducted in accordance with the guidelines outlined in the National Institutes of Health Guide for the Care and Use of Animals and were approved by the Institutional Animal Care and Use Committee at the University of Michigan.

#### 2.1.1 Testing schedule

After a one-week acclimation period, rats completed locomotor activity testing during week two and a five-minute elevated plus maze (EPM) session during week three. During week four, a subset of animals received tail-flick testing. Animals were euthanized for tissue collection (to obtain DNA for genotyping) during week five.

#### 2.1.2 Locomotor response to a novel environment

Locomotor response to a novel environment (hereafter, “locomotor”) was measured by placing each rat individually into a clean cage (23 cm × 43 cm) located in a different room with novel cues. The number of beam breaks (counts) was measured over 60 minutes for both lateral and rearing movements, defined as the Lateral and Rearing Locomotor Scores, respectively. The cumulative Lateral and Rearing Locomotor Scores over the 60-minute test were combined to generate the Total Locomotor Score. More details on the testing apparatus can be found in Stead et al, 2006.

#### 2.1.3 Elevated plus maze

Each rat was tested for five minutes on a standard EPM with two open arms and two closed arms. Video tracking quantified time allocation and movement (Ethovision, Noldus Information Technology). Details of the procedure are described in Chitre et al, 2023. We analyzed the following EPM traits: time in the open arms in seconds (EPM Open Arm Time), number of entries into the open arms (EPM Open Arm Entries), percent time in open arms across five minutes expressed as 0 to 100 percent (EPM Percent Time in Open Arms), total distance traveled in centimeters (EPM Distance Traveled), mean velocity in centimeters per second (EPM Velocity), and immobile time in seconds calculated as 300 minus moving time based on a center point criterion (EPM Time Immobile). The EPM Open Arm Time Ratio is the percent time in the open arms divided by the time in the open arms plus the closed arms. This ratio takes into account potential differences in the amount of time that animals spend in the center of the EPM. If center time is similar among animals, then the EPM Open Arm Time Ratio is comparable to the EPM Percent Time in Open Arms.

#### 2.1.4 Tail flick

Tail Flick Latency was recorded in week four. To assess individual differences in baseline pain thresholds to a noxious thermal stimulus, we used a modified tail flick (thermal nociception) test. Rats were gently restrained, the distal 1.5 cm of the tail was exposed to thermal nociceptive input and the latency to flick out of the beam was recorded, with a maximum cutoff time of 15s to prevent tissue damage. While we calibrated the test to produce an average latency of approximately 8s, we reduced the thermal intensity to increase the range of responses produced, allowing for better resolution of individual differences. Under lower thermal intensities, high reactivity rats demonstrate higher sensitivity to thermal stimulation (White et al, 2004), a difference that disappears at higher intensity stimulation. Each animal underwent two trials, separated by 90-120s, but due to inconsistent appearance of acute endogenous analgesia, we chose to only use the first trial in the analysis.

### 2.2 Statistical analysis

We first quantile normalized each trait. We then tested a set of candidate covariates that reflected the study design, including sex, age at the relevant behavioral test, test batch and apparatus identifiers for locomotor activity, experimenter and within-day test order for EPM and tail flick, and colony and housing room. For each trait, covariates that individually accounted for more than 2% of the phenotypic variance were retained, and the resulting covariate-adjusted values were used for all downstream analyses. Pairwise phenotypic correlations were estimated on the covariate-adjusted traits using Spearman’s ρ with two-sided p-values and all available data for each trait pair (pairwise complete observations); p ≤ 0.05 was considered significant. Sex was included as a candidate covariate for all traits and was retained (and regressed out) for several EPM measures but not for locomotor or tail flick traits; the retained covariate set for each trait is summarized in **Supplementary Table S2**.

### 2.3 Genotyping and Variant Processing

#### 2.3.1 DNA extraction, sequencing, and alignment

High molecular weight (HMW) genomic DNA from 654 outbred Sprague-Dawley spleen samples was isolated via the MagAttract HMW DNA Kit (Qiagen) at the University of Michigan and transferred to the University of California San Diego for further processing. All DNA samples were sequenced in low coverage using the Twist 96-Plex Library Preparation Kit (Twist Bioscience). Reads were demultiplexed with fgbio (http://fulcrumgenomics.github.io/fgbio/), trimmed with cutadapt (Martin, 2011), aligned to the GRCr8 reference genome with BWA-MEM (Li, 2013), and duplicate reads were removed with the Picard Toolkit (McKenna et al., 2010). Mean coverage was approximately 0.312x. In addition, eight of these 654 rats were deeply sequenced with KAPA HyperPre (Roche) followed by Illumina PE150 (Illumina) to approximately 19x; these eight outbred Sprague-Dawley samples formed a “truth set” to assess genotyping accuracy. Across these eight controls, the discordance between the final genotype data and the deep sequencing data was 0.99%, indicating high genotyping accuracy.

#### 2.3.2 Reference panel for imputation

We used inbred strains from the Hybrid Rat Diversity Panel (HRDP) as characterized in de Jong et al., 2024, to build the reference panel for STITCH-assisted imputation (Davies et al., 2016). Sequencing was available for 181 strains, of which 153 with the greatest depth were retained for the reference panel. Variants and genotypes for the reference panel were called with bcftools (Danecek, et al. 2021). Genotype posteriors below 0.99 were set to missing, only biallelic single-nucleotide polymorphisms (SNPs) with QUAL ≥ 20 and genotype missing rate ≤ 0.10 were retained. The resulting reference panel was then imputed and phased with Beagle 4.1 (*-gtgl* option) (Browning & Browning, 2016).

#### 2.3.3 Genotype calling and imputation

Genotypes for the 654 Sprague-Dawley rats were inferred with STITCH (reference-panel assisted). Sites supplied to STITCH included all loci present in the HRDP-derived reference panel and in the eight deeply sequenced Sprague-Dawley controls. We set *K* = 16 mosaic haplotypes and *n iterations* = 3. STITCH labeled genotypes with posterior probability < 0.90 were set as missing; missing genotypes were subsequently imputed with Beagle 4.1.

#### 2.3.4 Variant-level quality control

We began with 14,874,570 autosomal SNPs from the Beagle-imputed dataset. Variant filters were applied jointly on per-SNP genotype missingness (F-MISS), minor allele frequency (MAF), and Hardy-Weinberg equilibrium (HWE). We applied a slightly higher missingness threshold at loci that were present in either the HRDP-derived reference panel or the eight deeply sequenced Sprague-Dawley controls (STITCH “observed” sites; F-MISS ≤ 0.07) and a stricter threshold at loci supported only by imputation (imputed-only sites; F-MISS ≤ 0.04). In total, 1,643,596 autosomal SNPs passed all criteria and were retained for genetic analyses, and 13,230,974 SNPs were excluded by at least one criterion. Of the 9,088,280 SNPs with MAF < 0.05, 8,003,804 (88.1%) were monomorphic (MAF = 0) in this dataset. The distribution of variant failures across MAF, HWE, and missingness filters is summarized in **Supplementary Table S3**.

### 2.4 Genetic analysis

#### 2.4.1 Genetic relatedness, SNP heritability, and genetic correlation

We computed a genomic relatedness matrix (GRM) from the filtered autosomal SNP set using GCTA (*--make-grm*). SNP-based heritability for each trait was estimated via REML in GCTA. Pairwise genetic correlations (*r*_*g*_) were estimated using bivariate GREML. We performed exploratory bivariate GREML and, because many *r*_*g*_ estimates were boundary-constrained or had large standard errors, we did not report *r*_*g*_ values (Yang et al., 2011).

#### 2.4.2 GWAS and genome-wide thresholds

GWAS used GCTA linear mixed models (Yang et al., 2011) with a leave-one-chromosome-out GRM (LOCO) to reduce proximal contamination (Cheng and Palmer, 2013). Genome-wide significance thresholds were derived using permutation tests. The resulting thresholds were −log_10_*P* = 5.98 (α = 0.05) and −log_10_*P* = 5.60 (α = 0.10). Because all traits were quantile normalized, the same thresholds were used for all traits.

#### 2.4.3 QTL calling, conditional analysis, and variant annotation

For each locus, we defined the index SNP as the SNP with the smallest p-value (largest -log_10_*P*) within the region. We called a QTL when at least one SNP (the index SNP) exceeded the permutation-derived genome-wide threshold and a second SNP within 0.5 Mb had a p-value within 2 −log_10_*P* units of the index SNP. To identify additional independent loci on the same chromosome, we included the lead SNP(s) as covariates and iterated chromosome-specific scans until no further SNP exceeded the threshold. QTL intervals were defined as all markers in LD *r*^*2*^ >=0.6 with the peak SNP (Chitre et al., 2020). All variants within each QTL interval were annotated using SnpEff (Cingolani et al., 2012).

#### 2.4.4 Cross-study coordinate mapping and replication queries

The bHR/bLR F_2_ analyses used the Rnor_6.0 (RN6) reference, whereas the outbred Sprague-Dawley GWAS in this study used GRCr8. For phenotype-matched comparisons, we converted QTL peak SNPs and their LD-defined intervals (*r*^*2*^ ≥ 0.6) between builds using UCSC liftOver (Kuhn et al., 2013): outbred GRCr8 loci were lifted to RN6 for querying in the F_2_ summary statistics, and F_2_ RN6 loci were lifted to GRCr8 for querying in the outbred summary statistics. For each lifted locus we recorded the −log_10_*P* at the exact lifted top SNP when present and the maximum −log_10_*P* among all variants within the lifted LD-defined QTL interval. Because this procedure searches many correlated variants inside each interval, it is anti-conservative. Therefore, we treat these lookups as descriptive and do not regard them as confirmatory replication. We also ran an exploratory cross-phenotype interval-overlap screen.

## 3 Results

### 3.1 Phenotyping

All Sprague-Dawley rats were tested in three behavioral assays in the following order: locomotor activity in a novel environment, EPM, and tail flick. Measures from the first two assays contain exploratory locomotion traits with the EPM assay also measuring spontaneous anxiety-like behavior. Acute nociception was assessed in the tail flick test. **Figure 1** contains frequency distributions of these behaviors prior to quantile normalization of the data. Summary statistics for the trait data are reported in **Supplementary Table S4**. Apparent sex differences in the raw trait distributions (Figure 1) were modeled by including sex as a candidate covariate and regressing it out when it explained more than 2% of the phenotypic variance; the full set of covariates regressed out for each trait is reported in **Supplementary Table S2**.

**Figure 1.**
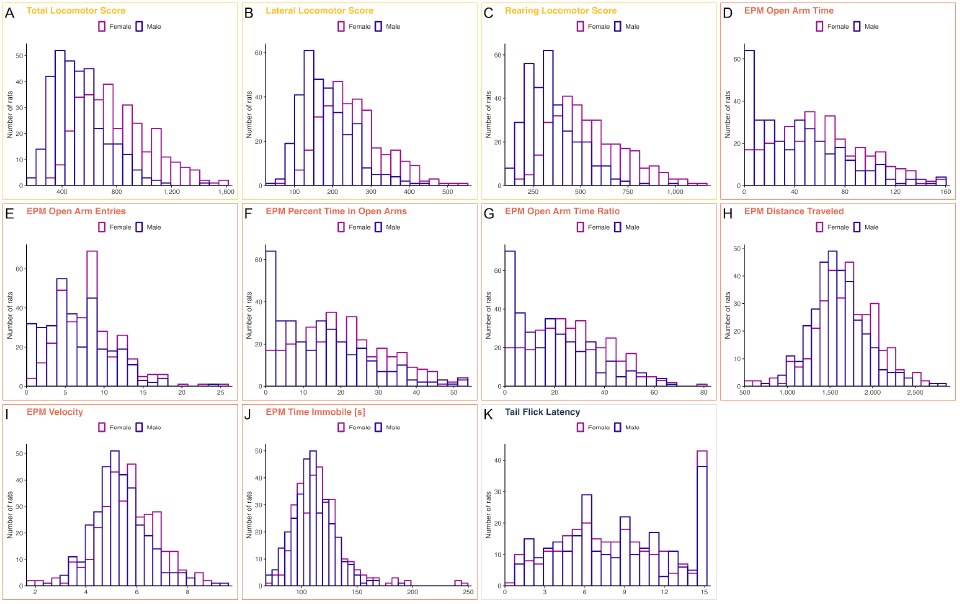
Frequency distributions of behavioral phenotypes by sex. Panels A–C (Locomotor response to a novel environment): Total Locomotor Score, Lateral Locomotor Score, Rearing Locomotor Score. Panels D–J (EPM): EPM Open Arm Time, EPM Open Arm Entries, EPM Percent Time in Open Arms, EPM Open Arm Time Ratio, EPM Distance Traveled, EPM Velocity, EPM Time Immobile. Panel K (Tail Flick): Tail Flick Latency. Data shown prior to quantile normalization. Summary statistics are in **Supplementary Table S4**; assay details and trait definitions are in Methods 2.1 (Phenotyping).

### 3.2 Phenotypic correlations

Several very high correlations were expected by design: EPM Distance Traveled and EPM Velocity are algebraically linked because velocity is distance over a fixed five-minute session (*ρ* ≈ 1.00); EPM Open Arm Time and EPM Percent Time in Open Arms are the same quantity rescaled by total time (*ρ* ≈ 1.00); and Total Locomotor Score aggregates Lateral Locomotor Score and Rearing Locomotor Score, explaining their high correlations (*ρ* ≈ 0.84-0.98). Within the EPM, EPM Percent Time in Open Arms and EPM Open Arm Time Ratio were highly correlated (*ρ* ≈ 0.96, *p* ≤ 0.05), and EPM Open Arm Entries correlated with both (*ρ* ≈ 0.71, *p* ≤ 0.05). EPM Distance Traveled and EPM Velocity are each related inversely to EPM Time Immobile (*ρ* ≈ −0.84, *p* ≤ 0.05). Tail Flick Latency showed limited associations, significant with two of 11 traits (*ρ* ≈ 0.10-0.14) and otherwise not significant. Results are shown in **Figure 2** and detailed values are provided in **Supplementary Table S5**.

**Figure 2.**
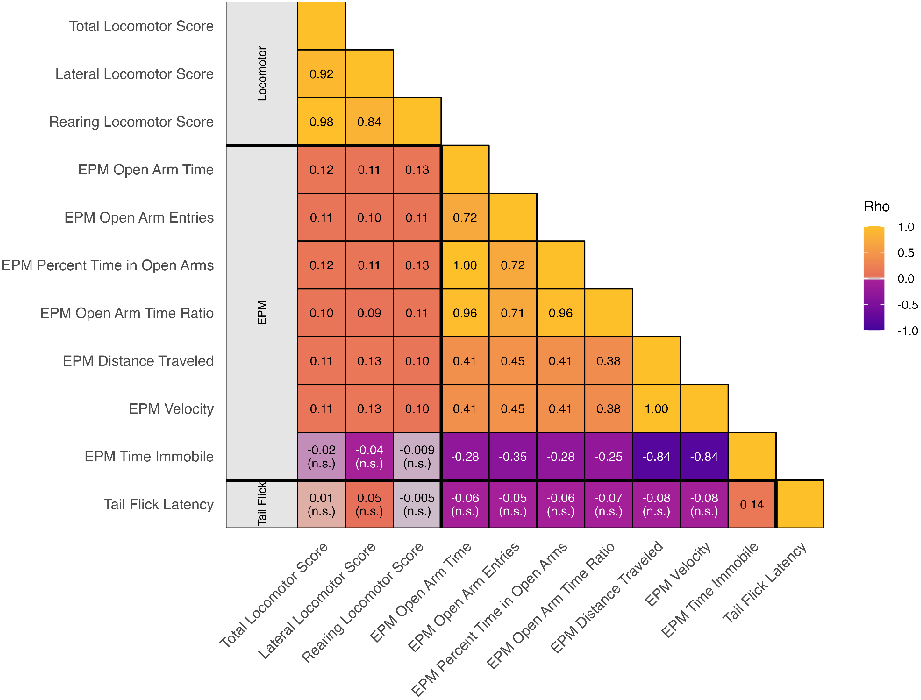
Phenotypic correlations among behavioral traits. Lower-triangular heatmap of Spearman rank correlations (Rho). Cells show Rho; entries labeled “n.s.” indicate p ≥ 0.05 (two-sided). Full Rho and p values are in **Supplementary Table S5**. Trait definitions are in Methods Section 2.1. Thick black lines delineate locomotor, EPM, and tail flick trait groups.

### 3.3 SNP heritability

Across 11 traits, SNP heritability (*h*^*2*^) ranged from 0.136 to 0.375. Locomotor traits showed the highest heritability estimates, exploratory locomotion and immobility measures in the EPM were moderate, and open-arm anxiety measures and tail flick latency were lower, with several open-arm measures not reaching statistical significance. Full values and p-values are provided in **Supplementary Table S6**.

### 3.4 GWAS

We identified two independent quantitative trait loci (QTLs) at the 5% genome-wide threshold and one additional QTL at the 10% threshold (-log_10_*P* = 5.98 at *α* = 0.05; -log_10_*P* = 5.60 at *α* = 0.10).

The strongest association was for EPM open-arm behavior on chromosome 1. The three open-arm EPM traits (EPM Percent Time in Open Arms, EPM Open Arm Time, EPM Open Arm Time Ratio) index the same construct and are simple rescalings, which is also reflected by their phenotypic correlations; accordingly, we treat them as a single locus on chromosome 1. At this locus, the top SNP for EPM Percent Time in Open Arms and EPM Open Arm Time was Chr1:267.773 Mb (−log_10_*P* = 7.28), whereas the top SNP for EPM Open Arm Time Ratio was Chr1:268.059 Mb (−log_10_*P* = 6.74); the LD interval at *r*^2^ ≥ 0.6 was 0.55 Mb. A moderate-impact missense variant in *Eno4* was identified within this interval (**Supplementary Table S7**). **Figure 3B** shows the regional association plot for EPM Percent Time in Open Arms at Chr1:267.773 Mb.

**Figure 3.**
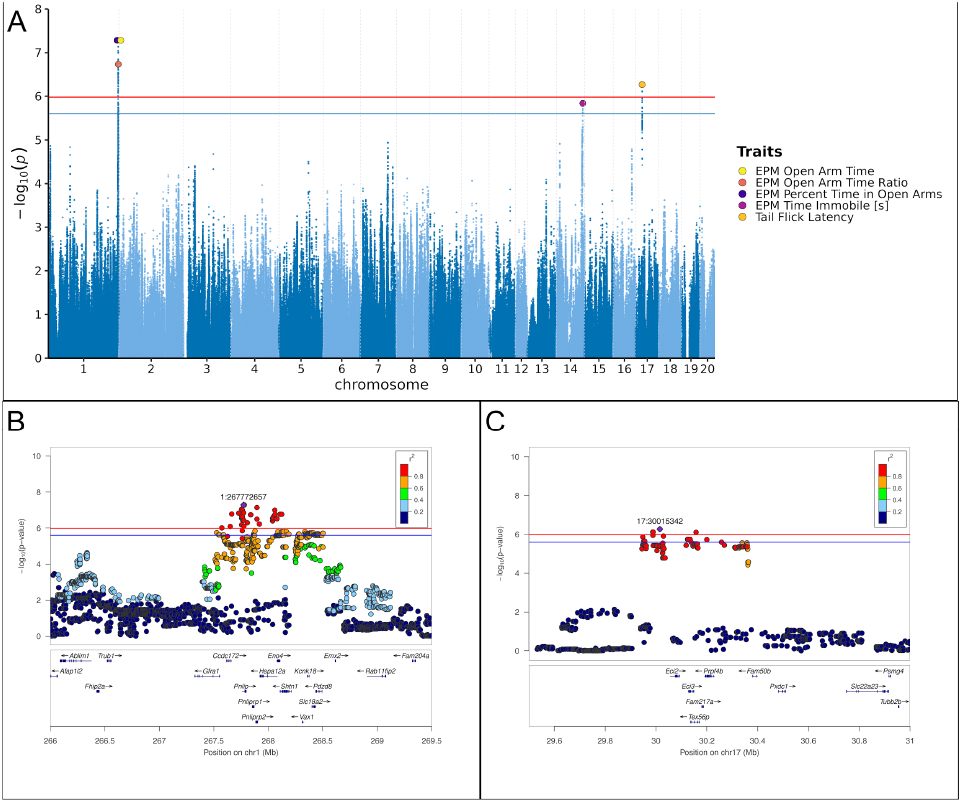
Genome-wide significant associations across behavioral traits, with representative loci at Chr1 (EPM Percent Time in Open Arms, 267.773 Mb) and Chr17 (Tail Flick Latency, 30.015 Mb) **A)** Combined Manhattan plot showing genome-wide significant hits. SNPs are plotted by genomic position (x) and -log_10_(*p*) (y) for traits with genome-wide significant associations. Red (genome-wide; -log_10_(*p*) = 5.98) and blue (suggestive; -log_10_(*p*) = 5.60) lines mark the significance thresholds. Large, filled markers highlight curated lead SNPs (colors = traits); overlapping peaks are slightly separated horizontally for visibility. **B)** Regional association plot for EPM Percent Time in Open Arms centered at the lead SNP (Chr1:267.773 Mb). Position (Mb) vs -log_10_(*p*); point color = LD (*r*^*2*^) to the lead SNP. Window 266.0-269.5 Mb; significance lines at 5.98 (genome-wide) and 5.60 (suggestive). Gene track displays genes across the region with arrows indicating transcription direction and labels by gene symbol. **C)** Regional association plot for Tail Flick Latency centered at the lead SNP (Chr17:30.015 Mb). Position (Mb) vs -log_10_(*p*); point color = LD (*r*^*2*^) to the lead SNP. Window 29.5-31.0 Mb; significance lines at 5.98 (genome-wide) and 5.60 (suggestive). Gene track displays genes across the region with arrows indicating transcription direction and labels by gene symbol.

Tail Flick Latency mapped to chromosome 17 with top SNP Chr17:30.015 Mb (−log_10_*P* = 6.27) and an LD interval of 0.41 Mb at *r*^2^ ≥ 0.6. A moderate-impact missense variant in *Tex56p* was identified within this interval (**Supplementary Table S7**). **Figure 3C** shows the regional association plot for Tail Flick Latency at Chr17:30.015 Mb.

EPM Time Immobile reached the 10% threshold on chromosome 14 with top SNP Chr14:102.124 Mb (−log_10_*P* = 5.84) and an LD interval of 0.13 Mb at *r*^2^ ≥ 0.6. No moderate-impact missense variants were observed in this interval.

**Figure 3A** shows the combined Manhattan plot for all genome-wide significant associations. Full statistics are provided in **Supplementary Table S8**. Manhattan plots for all traits are in **Supplementary Figure S1**. Additional regional association plots are in **Supplementary Figure S2**.

### 3.5 Comparison with bHR/bLR F_**2**_

#### 3.5.1 Phenotype set and age at testing

Here we compared the current GWAS in outbred Sprague-Dawley rats with our previous GWAS of an F_2_ intercross of selectively bred high-responder and low-responder (bHR/bLR) rats (Chitre et al., 2023). The studies used the same vivarium, apparatus, protocols, scoring rules, and preprocessing, and they assessed overlapping locomotor and EPM phenotypes. Age at testing differed between studies, with the F_2_ rats tested at older ages than the outbred Sprague-Dawley rats (**Supplementary Table S9**). Pavlovian Conditioned Approach (PavCA) was collected only in Chitre et al. (2023), whereas Tail Flick Latency was collected only in the current study.

#### 3.5.2 Direction of phenotypic correlations

In the bHR/bLR F_2_ rats, EPM Distance Traveled and EPM Percent Time in Open Arms were positively correlated with Lateral, Rearing, and Total Locomotor Scores, whereas EPM Time Immobile showed negative or weak correlations. In outbred Sprague-Dawley rats, the corresponding correlations were smaller and several were not significant despite the higher sample size in the present study. Side-by-side values for all overlapping pairs are provided in **Supplementary Table S10**.

#### 3.5.3 Cross-study QTL evaluation

In the bHR/bLR F_2_ rats, genome-wide significant loci were detected for Lateral Locomotor Score, Total Locomotor Score, Rearing Locomotor Score, and EPM Distance Traveled (Chitre et al., 2023). For the same phenotypes measured in both studies, we lifted outbred Sprague-Dawley GRCr8 QTLs to RN6 and queried them in the F_2_summary statistics, and reciprocally lifted F_2_ RN6 QTLs to GRCr8 and queried them in the outbred Sprague-Dawley summary statistics. We then performed the same liftover-based comparison for cross-phenotype pairs (e.g., F_2_ locomotor intervals vs. outbred EPM intervals). In all cases, there were no overlapping LD-defined QTL intervals between the two studies; when signals occurred on the same chromosome, peaks were separated by more than 100 Mb. Several lifted intervals contained variants with nominal p < 0.05 in the reciprocal dataset, but these represent the minimum p selected from many correlated tests within each interval and are therefore anti-conservative. We report them for completeness in **Supplementary Tables S11–S12** and do not interpret them as evidence of replication.

## 4 Discussion

In this study we performed a GWAS in commercially outbred Charles River Sprague-Dawley rats tested in young adulthood across three paradigms: locomotor response to a novel environment, EPM, and tail flick test. We analyzed 11 phenotypes. We chose these paradigms for their relevance to substance use liability and maintenance, with novelty-related traits forecasting acquisition, escalation, and reinstatement of psychostimulant self-administration (Flagel et al, 2014; Piazza et al., 1989; Gancarz et al., 2023), approach versus avoidance on the EPM relating to motivation for cocaine intake and supported by mixed-reality human analogs (Bush and Vaccarino 2007; Dilleen et al., 2012; Biedermann et al., 2017), and thermal nociception and pain states modulating opioid seeking and consumption (Hipólito et al., 2015; Koob, 2021; Higginbotham et al., 2022).

In a separate study, we previously performed a GWAS in a bHR x bLR F_2_intercross derived from Sprague-Dawley rats (Chitre et al., 2023). However, we did not find any significant overlap between the findings across the two studies, as the two populations appear to have captured different facets of addiction-relevant traits. While most of the traits for which loci were identified in the selectively bred F_2_ rats reflect facets of exploratory locomotion in a novel environment, most of the traits identified in the outbred Sprague-Dawley reflect loci for anxiety-related traits and not exploration. This difference in significant loci may reflect insufficient power, different alleles or allele frequencies between the two populations (perhaps due to selection for the HR and LR phenotypes), or age differences between the two cohorts or other factors. Our findings shed light on genetic differences associated with comparable exploratory locomotion and anxiety-like behaviors in outbred Sprague-Dawley rats vs. selectively bred rats on the same background.

### 4.1 Phenotypic correlations align with assay expectations

Several very high correlations were expected by design: EPM Distance Traveled and EPM Velocity, indexes of general activity, are algebraically linked because velocity is distance over a fixed five-minute session, EPM Open Arm Time and EPM Percent Time in Open Arms are the same quantity rescaled by total time, and Total Locomotor Score aggregates Lateral Locomotor Score and Rearing Locomotor Score. Among anxiety-related measures, EPM Open Arm Time and EPM Open Arm Entries were highly positively correlated and were each inversely correlated with EPM Time Immobile. EPM Time Immobile was also inversely correlated with the EPM general activity measures (i.e., Distance Traveled and EPM Velocity). This pattern matches standard interpretations of the EPM for anxiety-like behavior and locomotor activity within the apparatus (Walf & Frye, 2007).

Cross-assay correlations for distance were weak (for example, EPM Distance Traveled vs Lateral Locomotor Score *ρ* ≈ 0.11). This indicates that EPM movement is recorded under risk and immobility tendencies in outbred Sprague-Dawley rats, so it shares little variance with free locomotion in a walled novel cage (La-Vu et al., 2020; Gaspar et al., 2023; Snyder et al., 2021). In contrast, moderate cross-assay correlations for distance were found in the bHR/bLR F_2_ rats (Chitre et al, 2023), highlighting differences in the outbred vs selectively bred (for a propensity to explore a novel environment) rat lines. Tail Flick Latency also showed minimal correlations with EPM and locomotor measures in the outbreds aside from a small positive association with EPM Time Immobile. The sample size was sufficient; the pattern likely stems from higher measurement noise in tail flick latency (EPM ≈ 654, locomotor ≈ 650, tail flick 534).

### 4.2 SNP heritability of behavioral traits

Across 11 traits, SNP heritability estimates ranged from 0.136 to 0.375. Estimates were highest for the locomotor traits, moderate for EPM movement indices, modest for tail flick latency, and lowest for EPM open-arm metrics; three open-arm estimates did not reach *p* < 0.05. Although lower overall, this range of SNP heritability estimates from higher for exploratory locomotion traits to lower for anxiety-related traits is consistent with that found in our previous study examining bHR/bLR F_2_ rats (Chitre et al, 2023). The current results indicate that common variants account for a measurable share of trait variance in this outbred Sprague-Dawley population, supporting its use for GWAS (Gileta et al., 2022).

### 4.3 GWAS results

We studied young adult outbred Sprague Dawley rats across three paradigms capturing facets of exploratory locomotion, anxiety-like behavior, and nociception and conducted GWAS on 11 behavioral phenotypes. Although locomotor traits showed the highest SNP heritability, no significant QTLs were identified for them. We identified three independent QTLs for other traits, two for anxiety-related traits and one for nociception. The three EPM open arm measures, which quantify the same underlying construct (see Methods, Phenotyping), mapped to a single chromosome 1 locus, EPM Time Immobile mapped to chromosome 14, and Tail Flick Latency mapped to chromosome 17. Across traits, sample sizes ranged from 534 to 654 rats, which reduces statistical power and is a limitation of the study. In the subsections below we summarize each interval and nominate positional candidates.

#### 4.3.1 Chromosome 1 locus for open-arm exploration in the EPM

Multiple genes in the chromosome 1 interval provided plausible mechanisms for individual differences in anxiety-related open-arm behavior. *Slc18a2* encodes VMAT2, the vesicular monoamine transporter that sets vesicular loading of dopamine, norepinephrine, and serotonin. VMAT2 gain-of-function increases dopamine release and improves emotional behavior in mice, consistent with a monoaminergic pathway for EPM variation (Lohr et al. 2014; Branco et al. 2020). *Gfra1*, the co-receptor for GDNF, is required for mature synaptic connectivity in the medial habenula, a node that links limbic forebrain inputs to monoaminergic midbrain outputs and shapes anxiety and fear behaviors (Cacalano et al. 1998; Fernández-Suárez et al. 2021). *Pdzd8* regulates endoplasmic reticulum-mitochondria tethering and dendritic calcium handling, and its disruption in mice produces abnormalities in exploration and anxiety-relevant behaviors, providing a non-monoaminergic route to altered approach-avoidance (Hirabayashi et al. 2017; Kurihara et al. 2023). We identified a moderate-impact missense variant in *Eno4*, but *Eno4* is not expressed in brain (it is largely testis-specific), which makes it a less likely driver of a CNS behavioral phenotype despite the coding change (Nakamura et al. 2013). Taken together, *Slc18a2, Gfra1*, and *Pdzd8* form a coherent mechanistic set spanning monoamine handling, habenular circuitry, and neuronal calcium signaling. Collectively, these candidates converge on monoaminergic and neuronal signaling pathways and motivate targeted follow-up in this locus.

#### 4.3.2 Chromosome 14 locus for EPM immobility

Immobility in the EPM indexes anxiety state and coping within a risk context. Within this interval on chromosome 14, transcriptional regulators emerge as candidates. *Rel* encodes c-Rel, an NF-κB subunit. NF-κB signaling influences affective behavior and hippocampal plasticity, and mouse work shows reduced anxiety-like behavior when the pathway is perturbed, supporting a role for *Rel* in exploratory suppression and immobility (Kassed and Herkenham 2004; Ahn et al. 2008). *Bcl11a* (CTIP1) controls cortical circuit development. Haploinsufficiency in humans and mice produces cognitive and behavioral alterations, consistent with forebrain mechanisms that could modulate immobility on the EPM (Dias et al. 2016). *Rel* and *Bcl11a* represent plausible positional candidates for this locus.

#### 4.3.3 Chromosome 17 locus for Tail flick latency

Tail flick latency reflects spinal and supraspinal processing of thermal nociception. The interval on chromosome 17 contains *Eci2* and *Eci3*, which encode enzymes involved in the β-oxidation of unsaturated fatty acids. Lipid-derived mediators are increasingly implicated in nociceptive processing, suggesting that regulatory variation in fatty-acid handling, particularly their oxidation, could alter pain sensitivity (Domenichiello et al., 2021; Shapiro et al, 2016). We also observed a moderate-impact missense variant in *Tex56p*, a testis-biased gene, but there are no known roles in pain. Given its expression pattern and lack of established links to nociception or pain, this missense variant is an unlikely causal driver. Overall, this locus appears novel and reinforces the established role of lipid metabolism in pain sensitivity.

### 4.4 Comparison with bHR/bLR F_2_

We sought to identify allelic variants associated with the same behavioral traits by performing a GWAS in outbred Sprague Dawley rats from Charles River and comparing the results with the bHR/bLR F_2_intercross rats (Chitre et al., 2023). In this comparison, no phenotype-matched genome-wide significant QTL intervals overlapped between the studies, and we did not observe cross-phenotype interval overlaps when F_2_ locomotor loci were compared with outbred EPM loci. After lifting discovery intervals between builds and querying the reciprocal summary statistics within the lifted LD-defined QTL intervals (*r*^*2*^ ≥ 0.6), several intervals showed nominal support (*p* < 0.05); however, these interval queries are anti-conservative because they select the minimum p-value across many correlated variants within each interval. We therefore treat these nominal signals as descriptive checks rather than evidence of replication.

Several factors could contribute to the lack of phenotype-matched replication. Limited statistical power is likely an important factor in both studies, given sample sizes and the polygenic architecture of these behavioral traits. In addition, allele frequencies and local LD likely differed between the bHR/bLR F_2_ and the outbred Sprague-Dawley cohort due to the history of selection in the bHR/bLR lineage and colony over time, which can influence which alleles are common enough to detect in GWAS (Gileta et al., 2022). Epistatic interactions could further limit replication if single-locus effects depend on genetic background (Phillips, 2008; Mackay, 2014; Sittig et al., 2016). Finally, the F_2_ rats were older at testing than the outbred rats (123 ± 5 days vs. 77.7 ± 0.9 days for locomotor and 85.2 ± 1.6 days for EPM), and adult-age differences can shift baseline novelty and anxiety-like behavior and alter apparent effect sizes (Imhof et al., 1993; Doremus et al., 2004). Nevertheless, both the outbred Sprague-Dawley and the bHR/bLR F_2_ GWASs provide valuable genetic insight into addiction-relevant behaviors observed in a normal distribution of the population compared to those generated through selective breeding based on a specific behavioral criterion for modeling individual differences and the extremes in human behavioral liabilities.

## 5 Conclusion

In conclusion, we found three independent QTLs in outbred Charles River Sprague-Dawley rats for anxiety-related traits in the EPM and nociception in the tail flick test. Candidate genes and annotations point to monoaminergic signaling, transcriptional regulation, and lipid metabolism. We compared these loci with those from an F_2_ intercross of selectively bred high-responder and low-responder rats (Chitre et al., 2023). We did not observe phenotype-matched replication. This comparison highlights genetic differences associated with comparable exploratory locomotion and anxiety-like behaviors in outbred Sprague-Dawley rats vs. selectively bred rats on the same background. Genetic characterization of both natural behavioral variation and that created by selective breeding provides insight into determining causal variants tied to individual differences in behavioral liabilities relevant to psychiatric and substance use disorders. Taken together, these results extend the genetic characterization of addiction-relevant behaviors in outbred Sprague-Dawley rats and motivate replication and targeted experimental tests of mechanism.

## 6 Acknowledgements

This work was supported by the NIDA Center for GWAS in Outbred Rats (P50DA037844), the Center for Genetics, Genomics, and Epigenetics of Substance Use Disorders in Outbred Rats (P30DA060810), NIDA U01DA043098 (HA, JL, AP), ONR 00014-19-1-2149 (HA), the Pritzker Neuropsychiatric Research Consortium (HA, SW) and the Hope for Depression Research Foundation (HDRF) (HA).

