## Supplementary Tables and Figures for "Genetic Architecture of Addiction-Relevant Behaviors in Outbred Sprague-Dawley Rats Reveals Loci for Anxiety-Like and Nociceptive Traits"

**Table S1. Demographic Information.** Behavioral assay, trait, sample size by sex, and testing age. Age values are mean ± SD in days at the time the assay was run. "M/F" gives the male and female counts. Within an assay block, all listed traits share the same sample size and age as the first row of that block unless otherwise noted. For trait definitions and procedures, see Methods Section 2.1.

**Table S2. Covariates Regressed Out for Each Trait After the >2% Variance Screen.** For each behavioral trait, the table lists covariates that individually explained more than 2% of the phenotypic variance and were therefore regressed out prior to downstream analyses. For each covariate, $R^2$ denotes the proportion of trait variance explained by that covariate alone, and Percent Variance Explained reports $100\times R^2$. Sex met the >2% criterion for several EPM measures but not for locomotor or tail flick traits.

**Table S3. Variant-Level Quality Control Summary.** Filter Category indicates which variant-level quality control filter(s) each SNP failed: minor allele frequency (MAF < 0.05), Hardy–Weinberg equilibrium (HWE $P < 1\times10^{-7}$), and/or per-SNP genotype missingness (F-MISS above the specified thresholds). Rows labeled MAF only, HWE only, and F-MISS only correspond to SNPs that failed only that single filter, whereas combination rows correspond to SNPs that failed multiple filters. Total SNPs gives the total number of autosomal SNPs before QC. As noted in the main text, of the 9,088,280 SNPs with MAF < 0.05, 8,003,804 (88.1%) were monomorphic (MAF = 0).

**Table S4. Summary Statistics for Behavioral Traits by Sex and Overall.** For each trait within each behavioral assay (Locomotor Response to a Novel Environment, Elevated Plus Maze, Tail Flick), columns report Mean ± SD, Median, IQR, Min, and Max for Female, Male, and Overall. Values are computed from data before quantile normalization. SD, standard deviation; IQR, interquartile range.

**Table S5. Phenotypic Correlations.** Columns: Trait 1, Trait 2, rho, p_value, sig. rho is the Spearman rank correlation coefficient; p_value is two-sided; sig indicates significance at $p \leq 0.05$ (yes/no). Correlations were computed on covariate-adjusted, quantile-normalized residuals using pairwise complete observations. Trait definitions are in Methods Section 2.1.

**Table S6**.**SNP-Based Heritability of Behavioral Traits.** Heritability ($h^2$) was estimated with GCTA GREML using a genomic relationship matrix. The p value column reports the GCTA GREML likelihood-ratio test p-value for $H_0$: h = 0. Values that appeared as 0.000 in source files are shown as <0.001. Traits are ordered by $h^2$ from highest to lowest. Trait definitions are provided in Methods Section 2.1.

**Table S7. Moderate-Impact Coding Variants within QTL Intervals.** Summary of coding variants in linkage disequilibrium (LD) with trait lead SNPs. Coordinates are on the GRCr8 rat genome build. Candidate Gene is the gene harboring the coding variant and does not imply causality. QTL: peak marker of the QTL. SNP in LD with Peak Marker: moderate-impact missense variant (per SnpEff) in LD ($r^2 \geq 0.6$) with the peak marker. LD $r^2$ with Trait Top SNP:

pairwise LD r² between the listed variant and the trait lead SNP, estimated in the outbred Sprague-Dawley sample. SNP Change: HGVS c. annotation. Amino Acid Change: HGVS p. annotation.

**Table S8**. **Summary of QTLs.** Peak markers and linkage-disequilibrium (LD) intervals for quantitative trait loci. Coordinates are on the GRCr8 rat genome build. $-\log_{10}P$ is the test statistic; Significance_Level indicates the 5% or 10% genome-wide threshold derived by permutation. A1 is the effect allele, A2 is the other allele, and Freq is the effect-allele frequency. b is the per-allele effect on the quantile-normalized trait with standard error se. The LD interval is defined as all variants with $r^2 \geq 0.6$ around the peak, reported as Start and Stop base-pair positions and physical Size in megabases.

**Table S9 Shared Phenotypes between the Outbred Sprague-Dawley (this study) and the bHR/bLR $F_2$ Rats (Chitre et al., 2023).** This table shows the mean age $\pm$ SD at the time of behavioral testing for the rats in the two studies.

**Table S10. Phenotypic Correlations (Spearman rho) between Elevated Plus Maze (EPM) Measures and Locomotor Traits in Outbred Sprague-Dawley (this study) and bHR/bLR $F_2$ rats (Chitre et al., 2023).** This table reports Spearman rho and two-sided p for each overlapping EPM × Locomotor trait pair measured in both studies. Values are shown as: rho_Outbred, p_Outbred, rho_F2, p_F2. Only the traits present in both studies appear in the rows of this table. Outbred Sprague-Dawley refers to this study; bHR/bLR $F_2$ refers to the intercross reported in Chitre et al., 2023.

**Table S11. bHR/bLR $F_2$ (RN6) QTLs lifted to GRCr8 and Queried in Outbred Sprague-Dawley.** This table lists QTLs discovered in the bHR/bLR $F_2$ study on RN6 and their lifted coordinates on GRCr8 used to query the outbred Sprague-Dawley GWAS. Columns: Behavioral Test, Trait ($F_2$ Label); Original Top SNP: $F_2$ peak SNP formatted as chr:pos; Original Top SNP $-\log_{10}P$: $F_2$ peak strength; Original LD Start: $F_2$ LD window start in bp; Original LD Stop: $F_2$ LD window stop in bp; Original Interval Size: $F_2$ LD window size in Mb computed as (stop − start)/1e6; Queried Trait: queried outbred trait; Lifted Top SNP: lifted peak SNP on GRCr8 formatted as chr:pos, NA if absent; Lifted LD Start: lifted LD window start in bp on GRCr8; Lifted LD Stop: lifted LD window stop in bp on GRCr8; Lifted Interval Size: lifted window size in Mb; Best $-\log_{10}P$: best signal in outbred Sprague-Dawleys within the lifted window reported as $-\log_{10}P$; Best SNP: position of that best signal formatted as chr:pos. Sizes are in megabases and rounded.

**Table S12. Outbred Sprague-Dawley (GRCr8) Loci Lifted to RN6 and Queried in bHR/bLR $F_2$.** This table lists QTLs discovered in outbred Sprague-Dawleys on GRCr8 and their lifted coordinates on RN6 used to query the bHR/bLR $F_2$ GWAS. Columns: Behavioral Test, Trait: outbred trait label; Original Top SNP: outbred peak SNP formatted as chr:pos; Original Top SNP -log10P: outbred peak strength; Original LD Start: outbred LD window start in bp; Original LD Stop: outbred LD window stop in bp; Original Interval Size (Mb): outbred window size in Mb; Queried Trait: queried $F_2$ trait; Lifted Top SNP: lifted peak SNP on RN6 formatted as chr:pos,

NA if absent; Lifted LD Start: lifted LD window start in bp on RN6; Lifted LD Stop: lifted LD window stop in bp on RN6; Lifted Interval Size (MB): lifted window size in Mb; Best −log10P: best signal in $F_2$ within the lifted window reported as −log10P; best SNP: position of that best signal formatted as chr:pos Sizes are in megabases and rounded.

**Table S1. Demographic Information.**

| Behavioral Assay | Trait | Sample Size (M/F) | Mean Age (Days) ± SD |
|---|---|---|---|
| Locomotor Response to a Novel Environment | Lateral Locomotor Score | 650 (327/323) | 77.68 ± 0.90 |
| | Total Locomotor Score | | |
| | Rearing Locomotor Score | | |
| Elevated Plus Maze | EPM Distance Traveled | 654 (327/327) | 85.22 ± 1.55 |
| | EPM Open Arm Entries | | |
| | EPM Percent Time in Open Arms | | |
| | EPM Open Arm Time Ratio | | |
| | EPM Open Arm Time | | |
| | EPM Velocity | | |
| | EPM Time Immobile | | |
| Tail Flick | Tail Flick Latency | 534 (267/267) | 94.03 ± 3.48 |

**Table S2. Covariates Regressed Out for Each Trait After the >2% Variance Screen.**

| Trait | Covariate Regressed Out | R2 | Percent Variance Explained |
|---|---|---|---|
| EPM Open Arm Time Ratio | Experimenter: EH_SK | 0.074 | 7.4 |
| EPM Open Arm Time Ratio | Sex: M | 0.071 | 7.1 |
| EPM Open Arm Time Ratio | Experimenter: EH_KA | 0.02 | 2 |
| EPM Open Arm Time | Experimenter: EH_SK | 0.092 | 9.2 |
| EPM Open Arm Time | Sex: M | 0.065 | 6.5 |
| EPM Open Arm Time | Experimenter: EH_JS | 0.028 | 2.8 |
| EPM Open Arm Time | Experimenter: EH_KA | 0.027 | 2.7 |
| EPM Open Arm Entries | Sex: M | 0.053 | 5.3 |
| EPM Open Arm Entries | Experimenter: EH_SM | 0.033 | 3.3 |
| EPM Open Arm Entries | room: R04 or R08 | 0.032 | 3.2 |
| EPM Open Arm Entries | Experimenter: EH_JS | 0.022 | 2.2 |
| EPM Percent Time in Open Arms | Experimenter: EH_SK | 0.092 | 9.2 |
| EPM Percent Time in Open Arms | Sex: M | 0.065 | 6.5 |
| EPM Percent Time in Open Arms | Experimenter: EH_JS | 0.028 | 2.8 |
| EPM Percent Time in Open Arms | Experimenter: EH_KA | 0.027 | 2.7 |
| EPM Distance Traveled | Experimenter: EH_SM | 0.04 | 4 |
| EPM Distance Traveled | room: R04 or R08 | 0.038 | 3.8 |
| EPM Distance Traveled | Sex: M | 0.02 | 2 |
| EPM Velocity | Experimenter: EH_SM | 0.04 | 4 |
| EPM Velocity | room: R04 or R08 | 0.038 | 3.8 |
| EPM Velocity | Sex: M | 0.02 | 2 |
| EPM Time Immobile | Experimenter: EH_JS | 0.031 | 3.1 |
| EPM Time Immobile | Test Order Per Day: 29 | 0.02 | 2 |

**Table S3. Variant-Level Quality Control Summary.**

| Filter Category | Count |
|---|---|
| MAF only | 8,042,158 |
| HWE only | 1,460 |
| F-MISS only | 3,911,924 |
| MAF+HWE | 2,415 |
| MAF+F-MISS | 1,041,835 |
| HWE+F-MISS | 229,310 |
| MAF+HWE+F-MISS | 1,872 |
| Total SNPs | 14,874,570 |

**Table S4. Summary Statistics for Behavioral Traits by Sex and Overall.**

| Behavioral Assay | Trait | Female | | | | | Male | | | | | Overall | | | | |
|---|---|---|---|---|---|---|---|---|---|---|---|---|---|---|---|---|
| | | Mean ± SD | Median | IQR | Min | Max | Mean ± SD | Median | IQR | Min | Max | Mean ± SD | Median | IQR | Min | Max |
| Locomotor Response to a Novel Environment | Total Locomotor Score | 794.57 ± 259.14 | 746.00 | 369.50 | 301.00 | 1,577.00 | 527.89 ± 192.59 | 498.00 | 248.00 | 172.00 | 1,413.00 | 660.41 ± 264.11 | 608.00 | 350.50 | 172.00 | 1,577.00 |
| | Lateral Locomotor Score | 251.45 ± 81.86 | 239.00 | 107.00 | 58.00 | 550.00 | 181.91 ± 66.78 | 170.00 | 91.00 | 42.00 | 429.00 | 216.47 ± 82.31 | 204.00 | 108.75 | 42.00 | 550.00 |
| | Rearing Locomotor Score | 543.12 ± 189.13 | 510.00 | 264.00 | 172.00 | 1,119.00 | 345.98 ± 135.52 | 325.00 | 173.50 | 104.00 | 987.00 | 443.94 ± 191.58 | 412.00 | 245.75 | 104.00 | 1,119.00 |
| Elevated Plus Maze | EPM Open Arm Time | 61.30 ± 35.58 | 58.03 | 52.15 | 0.00 | 160.50 | 43.75 ± 36.59 | 39.30 | 52.62 | 0.00 | 160.26 | 52.53 ± 37.12 | 48.78 | 53.39 | 0.00 | 160.50 |
| | EPM Open Arm Entries | 8.10 ± 3.94 | 8.00 | 5.00 | 0.00 | 26.00 | 6.46 ± 4.15 | 6.00 | 6.00 | 0.00 | 24.00 | 7.28 ± 4.13 | 7.00 | 6.00 | 0.00 | 26.00 |
| | EPM Percent Time in Open Arms | 20.50 ± 11.90 | 19.41 | 17.44 | 0.00 | 53.68 | 14.63 ± 12.24 | 13.14 | 17.60 | 0.00 | 53.60 | 17.57 ± 12.41 | 16.32 | 17.86 | 0.00 | 53.68 |
| | EPM Open Arm Time Ratio | 27.20 ± 15.56 | 25.96 | 23.01 | 0.00 | 81.22 | 19.07 ± 15.81 | 17.18 | 23.29 | 0.00 | 66.09 | 23.13 ± 16.19 | 21.66 | 24.38 | 0.00 | 81.22 |
| | EPM Distance Traveled | 1,683.70 ± 353.29 | 1,690.33 | 454.93 | 522.42 | 2,620.64 | 1,592.21 ± 310.72 | 1,567.28 | 356.44 | 725.40 | 2,825.23 | 1,637.96 ± 335.57 | 1,621.08 | 428.46 | 522.42 | 2,825.23 |
| | EPM Velocity | 5.63 ± 1.18 | 5.66 | 1.52 | 1.75 | 8.77 | 5.33 ± 1.04 | 5.25 | 1.19 | 2.43 | 9.46 | 5.48 ± 1.12 | 5.42 | 1.44 | 1.75 | 9.46 |
| | EPM Time Immobile | 116.95 ± 23.78 | 114.70 | 25.32 | 70.50 | 246.83 | 110.65 ± 18.39 | 108.83 | 22.52 | 70.53 | 188.20 | 113.80 ± 21.47 | 111.50 | 24.61 | 70.50 | 246.83 |
| Tail Flick | Tail Flick Latency | 8.42 ± 4.24 | 8.08 | 6.56 | 0.52 | 15.00 | 8.24 ± 4.26 | 7.96 | 6.39 | 0.88 | 15.00 | 8.33 ± 4.24 | 8.00 | 6.40 | 0.52 | 15.00 |

**Table S5. Phenotypic Correlations.**

| Trait 1 | Trait 2 | rho | p_value | sig |
|---|---|---|---|---|
| Lateral Locomotor Score | Total Locomotor Score | 0.92 | 3.40E-265 | yes |
| Rearing Locomotor Score | Total Locomotor Score | 0.98 | 0.00E+00 | yes |
| Rearing Locomotor Score | Lateral Locomotor Score | 0.84 | 1.80E-174 | yes |
| EPM Open Arm Time | Total Locomotor Score | 0.12 | 1.47E-03 | yes |
| EPM Open Arm Time | Lateral Locomotor Score | 0.11 | 6.18E-03 | yes |
| EPM Open Arm Time | Rearing Locomotor Score | 0.13 | 1.05E-03 | yes |
| EPM Open Arm Entries | Total Locomotor Score | 0.11 | 5.50E-03 | yes |
| EPM Open Arm Entries | Lateral Locomotor Score | 0.1 | 8.96E-03 | yes |
| EPM Open Arm Entries | Rearing Locomotor Score | 0.11 | 5.37E-03 | yes |
| EPM Open Arm Entries | EPM Open Arm Time | 0.72 | 5.60E-105 | yes |
| EPM Percent Time in Open Arms | Total Locomotor Score | 0.12 | 1.47E-03 | yes |
| EPM Percent Time in Open Arms | Lateral Locomotor Score | 0.11 | 6.18E-03 | yes |
| EPM Percent Time in Open Arms | Rearing Locomotor Score | 0.13 | 1.05E-03 | yes |
| EPM Percent Time in Open Arms | EPM Open Arm Time | 1 | 0.00E+00 | yes |
| EPM Percent Time in Open Arms | EPM Open Arm Entries | 0.72 | 5.60E-105 | yes |
| EPM Open Arm Time Ratio | Total Locomotor Score | 0.1 | 9.08E-03 | yes |
| EPM Open Arm Time Ratio | Lateral Locomotor Score | 0.09 | 2.95E-02 | yes |
| EPM Open Arm Time Ratio | Rearing Locomotor Score | 0.11 | 6.45E-03 | yes |
| EPM Open Arm Time Ratio | EPM Open Arm Time | 0.96 | 0.00E+00 | yes |
| EPM Open Arm Time Ratio | EPM Open Arm Entries | 0.71 | 4.30E-103 | yes |
| EPM Open Arm Time Ratio | EPM Percent Time in Open Arms | 0.96 | 0.00E+00 | yes |
| EPM Distance Traveled | Total Locomotor Score | 0.11 | 3.68E-03 | yes |
| EPM Distance Traveled | Lateral Locomotor Score | 0.13 | 6.11E-04 | yes |
| EPM Distance Traveled | Rearing Locomotor Score | 0.1 | 8.61E-03 | yes |
| EPM Distance Traveled | EPM Open Arm Time | 0.41 | 7.00E-28 | yes |
| EPM Distance Traveled | EPM Open Arm Entries | 0.45 | 2.56E-33 | yes |
| EPM Distance Traveled | EPM Percent Time in Open Arms | 0.41 | 7.00E-28 | yes |
| EPM Distance Traveled | EPM Open Arm Time Ratio | 0.38 | 9.40E-24 | yes |
| EPM Velocity | Total Locomotor Score | 0.11 | 0.00373 | yes |

| | | | | |
|---|---|---|---|---|
| EPM Velocity | Lateral Locomotor Score | 0.13 | 0.000632 | yes |
| EPM Velocity | Rearing Locomotor Score | 0.1 | 0.00867 | yes |
| EPM Velocity | EPM Open Arm Time | 0.41 | 6.38E-28 | yes |
| EPM Velocity | EPM Open Arm Entries | 0.45 | 2.46E-33 | yes |
| EPM Velocity | EPM Percent Time in Open Arms | 0.41 | 6.38E-28 | yes |
| EPM Velocity | EPM Open Arm Time Ratio | 0.38 | 8.53E-24 | yes |
| EPM Velocity | EPM Distance Traveled | 1 | 0 | yes |
| EPM Time Immobile | Total Locomotor Score | -0.02 | 0.665 | no |
| EPM Time Immobile | Lateral Locomotor Score | -0.04 | 0.278 | no |
| EPM Time Immobile | Rearing Locomotor Score | -0.01 | 0.826 | no |
| EPM Time Immobile | EPM Open Arm Time | -0.28 | 1.20E-13 | yes |
| EPM Time Immobile | EPM Open Arm Entries | -0.35 | 2.06E-20 | yes |
| EPM Time Immobile | EPM Percent Time in Open Arms | -0.28 | 1.20E-13 | yes |
| EPM Time Immobile | EPM Open Arm Time Ratio | -0.25 | 4.47E-11 | yes |
| EPM Time Immobile | EPM Distance Traveled | -0.84 | 4.00E-177 | yes |
| EPM Time Immobile | EPM Velocity | -0.84 | 2.90E-177 | yes |
| Tail Flick Latency | Total Locomotor Score | 0.01 | 0.739 | no |
| Tail Flick Latency | Lateral Locomotor Score | 0.05 | 0.247 | no |
| Tail Flick Latency | Rearing Locomotor Score | -0.01 | 0.907 | no |
| Tail Flick Latency | EPM Open Arm Time | -0.06 | 0.188 | no |
| Tail Flick Latency | EPM Open Arm Entries | -0.05 | 0.283 | no |
| Tail Flick Latency | EPM Percent Time in Open Arms | -0.06 | 0.188 | no |
| Tail Flick Latency | EPM Open Arm Time Ratio | -0.07 | 0.12 | no |
| Tail Flick Latency | EPM Distance Traveled | -0.08 | 0.0742 | no |
| Tail Flick Latency | EPM Velocity | -0.08 | 0.0727 | no |
| Tail Flick Latency | EPM Time Immobile | 0.14 | 0.00128 | yes |

**Table S6.SNP-Based Heritability of Behavioral Traits.**

| Trait | SNP h2 ± SE | P value |
|---|---|---|
| Lateral Locomotor Score | 0.375 ± 0.083 | <0.001 |
| Total Locomotor Score | 0.368 ± 0.086 | <0.001 |
| Rearing Locomotor Score | 0.351 ± 0.087 | <0.001 |
| EPM Velocity | 0.298 ± 0.086 | <0.001 |
| EPM Distance Traveled | 0.296 ± 0.086 | <0.001 |
| EPM Time Immobile | 0.276 ± 0.084 | <0.001 |
| EPM Open Arm Time Ratio | 0.156 ± 0.082 | 0.028 |
| Tail Flick Latency | 0.153 ± 0.094 | 0.033 |
| EPM Percent Time in Open Arms | 0.143 ± 0.083 | 0.054 |
| EPM Open Arm Time | 0.143 ± 0.083 | 0.054 |
| EPM Open Arm Entries | 0.136 ± 0.084 | 0.061 |

**Table S7. Moderate-Impact Coding Variants within QTL Intervals.**

| Trait | QTL | Candidate Gene | SNP in LD with Peak Marker | LD $r^2$ with Trait Top SNP | SNP Change | Amino Acid Change |
|---|---|---|---|---|---|---|
| EPM Percent Time in Open Arms | Chr1:267.773 Mb | *Eno4* | Chr1:268,084,478 | 0.681 | c.187G>T | p.Ala63Ser |
| EPM Open Arm Time Ratio | Chr1:268.059 Mb | *Eno4* | Chr1:268,084,478 | 0.691 | c.187G>T | p.Ala63Ser |
| EPM Open Arm Time | Chr1:267.773 Mb | *Eno4* | Chr1:268,084,478 | 0.681 | c.187G>T | p.Ala63Ser |
| Tail Flick Latency | Chr17:30.015 Mb | *Tex56p* | Chr17:30,152,157 | 0.839 | c.552C>G | p.Asp184Glu |

**Table S8. Summary of QTLs.**

| | Peak Marker | | | | | | | | | | LD Interval | | |
|---|---|---|---|---|---|---|---|---|---|---|---|---|---|
| Trait | Chr | SNP | Position (bp) | -log10P | Significance_Level | A1 | A2 | Freq | b | se | Start (bp) | Stop (bp) | Size (Mb) |
| EPM Percent Time in Open Arms | Chr1 | Chr1:267.773 Mb | 267,772,657 | 7.283 | 5% | A | T | 0.453 | -0.314 | 0.058 | 267,526,572 | 268,502,952 | 0.55 |
| EPM Open Arm Time | | Chr1:267.773 Mb | 267,772,657 | 7.283 | 5% | A | T | 0.453 | -0.314 | 0.058 | 267,526,572 | 268,502,952 | 0.55 |
| EPM Open Arm Time Ratio | | Chr1:268.059 Mb | 268,059,169 | 6.735 | 5% | G | A | 0.455 | -0.303 | 0.058 | 267,527,231 | 268,486,673 | 0.55 |
| EPM Time Immobile | Chr14 | Chr14:102.124 Mb | 102,123,644 | 5.838 | 10% | T | C | 0.498 | 0.272 | 0.057 | 101,958,495 | 102,170,686 | 0.13 |
| Tail Flick Latency | Chr17 | Chr17:30.015 Mb | 30,015,342 | 6.267 | 5% | A | T | 0.089 | -0.546 | 0.109 | 29,944,792 | 30,363,871 | 0.41 |

**Table S9. Shared Phenotypes between the Outbred Sprague-Dawley (this study) and the bHR/bLR $F_2$ Rats (Chitre et al., 2023).**

| Assay | Shared Phenotype | Outbred Age (days; mean ± SD) | F2 Age (days; mean ± SD) | Δ Age (weeks) |
|---|---|---|---|---|
| Locomotor response to novelty | Lateral Locomotor Score | 77.7 ± 0.9 | 123 ± 5 | ~6.5 |
| | Rearing Locomotor Score | | | |
| | Total Locomotor Score | | | |
| Elevated plus maze | EPM Distance Traveled | 85.2 ± 1.6 | 123 ± 5 | ~5.4 |
| | EPM Percent Time in Open Arms | | | |
| | EPM Time Immobile | | | |

**Table S10. Phenotypic Correlations (Spearman rho) between Elevated Plus Maze (EPM) Measures and Locomotor Traits in Outbred Sprague-Dawley (this study) and bHR/bLR $F_2$ rats (Chitre et al., 2023).**

| EPM Measure | Locomotor Measure | rho_Outbred | p_Outbred | rho_F2 | p_F2 |
|---|---|---|---|---|---|
| EPM Distance Traveled | Lateral Locomotor Score | 0.13 | 1.00E-03 | 0.402 | 0.00E+00 |
| EPM Distance Traveled | Total Locomotor Score | 0.11 | 4.00E-03 | 0.424 | 0.00E+00 |
| EPM Distance Traveled | Rearing Locomotor Score | 0.1 | 9.00E-03 | 0.414 | 0.00E+00 |
| EPM Percent Time in Open Arms | Lateral Locomotor Score | 0.11 | 6.00E-03 | 0.244 | 5.39E-07 |
| EPM Percent Time in Open Arms | Total Locomotor Score | 0.12 | 1.00E-03 | 0.262 | 7.22E-08 |
| EPM Percent Time in Open Arms | Rearing Locomotor Score | 0.13 | 1.00E-03 | 0.255 | 1.70E-07 |
| EPM Time Immobile | Lateral Locomotor Score | -0.04 | 2.78E-01 | -0.358 | 7.80E-14 |
| EPM Time Immobile | Total Locomotor Score | -0.02 | 6.65E-01 | -0.417 | 0.00E+00 |
| EPM Time Immobile | Rearing Locomotor Score | -0.01 | 8.26E-01 | -0.432 | 0.00E+00 |

**Table S11. bHR/bLR $F_2$ (RN6) QTLs lifted to GRCr8 and Queried in Outbred Sprague-Dawley.**

| Study & Trait | | Origin (bHR/bLR F2, RN6) | | | | | Lifted & Queried (Outbred Sprague-Dawley, GRCr8) | | | | | | |
|---|---|---|---|---|---|---|---|---|---|---|---|---|---|
| **Behavioral Test** | **Trait (F2 Label)** | **Original Top SNP** | **Original Top SNP -log10P** | **Original LD Start** | **Original LD Stop** | **Original Interval Size (Mb)** | **Queried Trait** | **Lifted Top SNP** | **Lifted LD Start** | **Lifted LD Stop** | **Lifted Interval Size (Mb)** | **Best −log10P** | **Best SNP** |
| Locomotor Response to a Novel Environment | Total Locomotor Score | Chr1:95,096,740 | 7.66 | 94,322,277 | 95,263,107 | 0.941 | Total Locomotor Score | NA | 99,762,095 | 100,692,809 | 0.931 | 1.61 | Chr1:99,782,419 |
| | Lateral Locomotor Score | Chr1:95,096,740 | 7.00 | 94,322,277 | 95,263,107 | 0.941 | Lateral Locomotor Score | NA | 99,762,095 | 100,692,809 | 0.931 | 1.30 | Chr1:99,782,419 |
| | Rearing Locomotor Score | Chr1:95,087,956 | 6.89 | 94,322,277 | 95,263,107 | 0.941 | Rearing Locomotor Score | NA | 99,762,095 | 100,692,809 | 0.931 | 1.66 | Chr1:99,862,586 |
| | Total Locomotor Score | Chr1:107,298,166 | 11.88 | 107,126,050 | 107,363,578 | 0.238 | Total Locomotor Score | NA | 110,477,039 | 110,718,512 | 0.241 | 1.45 | Chr1:110,485,783 |
| | Lateral Locomotor Score | Chr1:107,181,408 | 12.09 | 107,126,050 | 107,451,256 | 0.325 | Lateral Locomotor Score | NA | 110,477,039 | 110,806,642 | 0.33 | 1.12 | Chr1:110,485,783 |
| | Rearing Locomotor Score | Chr1:107,298,166 | 10.97 | 107,126,050 | 107,363,578 | 0.238 | Rearing Locomotor Score | NA | 110,477,039 | 110,718,512 | 0.241 | 1.42 | Chr1:110,485,783 |
| | Lateral Locomotor Score | Chr7:84,269,306 | 7.91 | 83,274,530 | 90,251,798 | 6.977 | Lateral Locomotor Score | NA | 77,406,383 | 83,965,203 | 6.559 | 2.26 | Chr7:80,705,907 |
| EPM | EPM Distance Traveled | Chr1:94,914,942 | 7.13 | 94,107,350 | 95,263,107 | 1.156 | EPM Distance Traveled | NA | 99,551,713 | 100,692,809 | 1.141 | 0.47 | Chr1:99,888,981 |

**Table S12. Outbred Sprague-Dawley (GRCr8) Loci Lifted to RN6 and Queried in bHR/bLR $F_2$.**

| Study & Trait | | Origin (Outbred Sprague-Dawley, GRCr8) | | | | | Lifted & Queried (bHR/bLR F2, RN6) | | | | | | |
|---|---|---|---|---|---|---|---|---|---|---|---|---|---|
| **Behavioral Test** | **Trait** | **Original Top SNP** | **Original Top SNP -log10P** | **Original LD Start** | **Original LD Stop** | **Original Interval Size (Mb)** | **Queried Trait** | **Lifted Top SNP** | **Lifted LD Start** | **Lifted LD Stop** | **Lifted Interval Size (MB)** | **Best −log10P** | **Best SNP** |
| EPM | Percent Time in Open Arms | Chr1:267,772,657 | 7.28 | 267,526,572 | 268,502,952 | 0.55 | EPM Percent Time in Open Arms | NA | 279,542,985 | 280,525,689 | 0.98 | 2.23 | Chr1:280,052,026 |
| | Time Immobile | Chr14:102,123,644 | 5.84 | 101,958,495 | 102,170,686 | 0.13 | EPM Time Immobile | NA | 108,554,098 | 108,766,958 | 0.21 | 1.62 | Chr14:108,679,940 |

**Supplementary Figure S1.** Manhattan plots for each behavioral trait. SNPs are plotted by genomic position on GRCr8 (x) and −log10P from LOCO mixed-model GWAS (y). Horizontal lines show genome-wide (5% at −log10P = 5.98) and suggestive (10% at −log10P = 5.60) thresholds derived by permutation.

EPM Distance Traveled

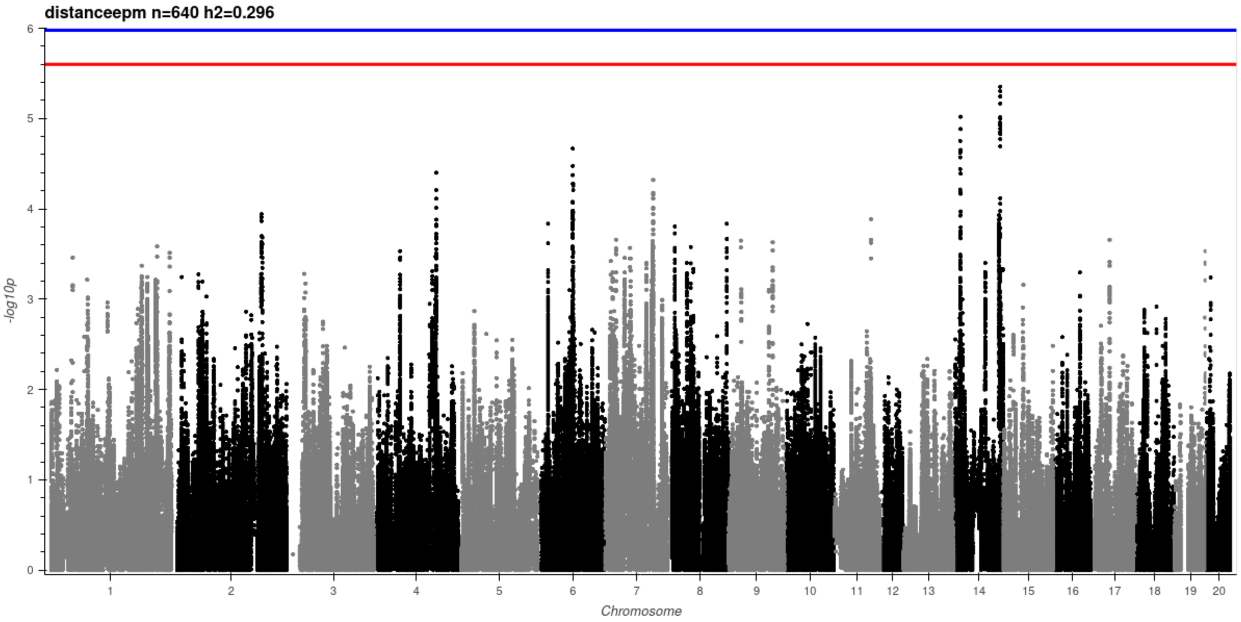


EPM Open Arm Entries

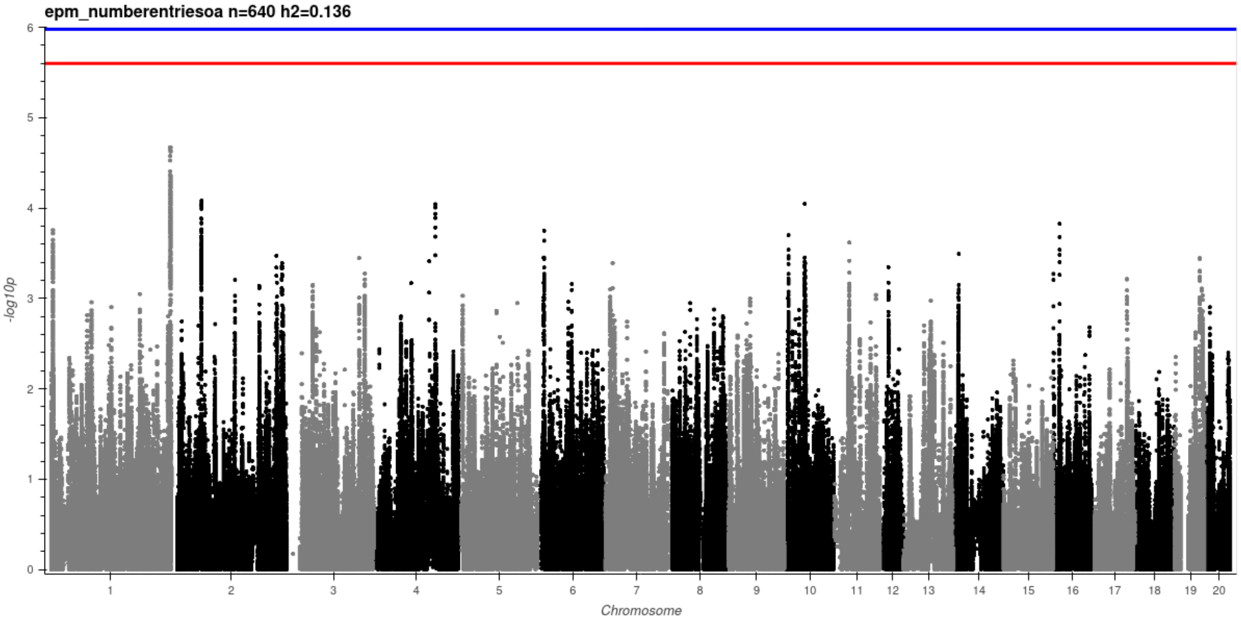

### EPM Percent Time in Open Arms

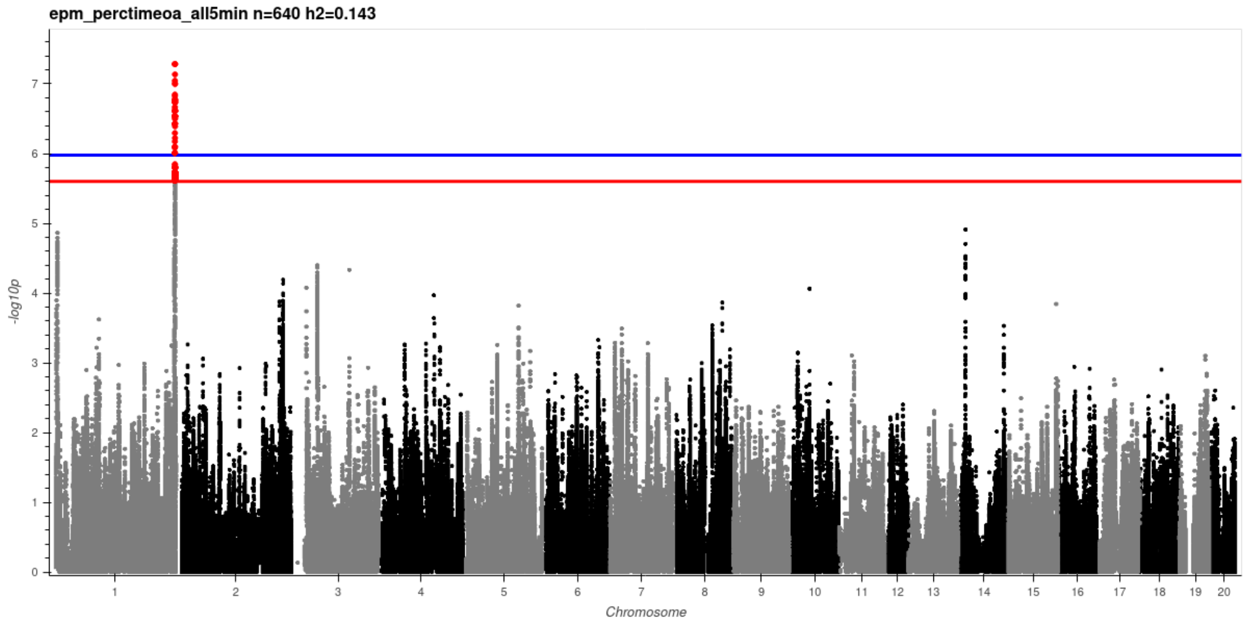


### EPM Open Arm Time Ratio

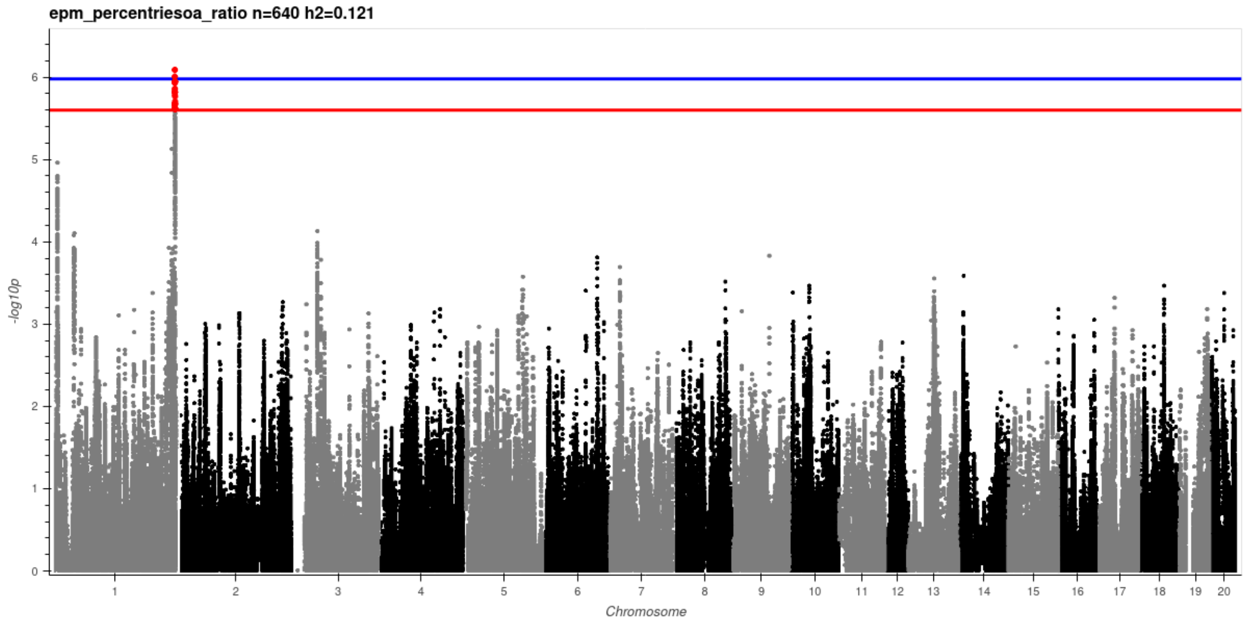

EPM Open Arm Time

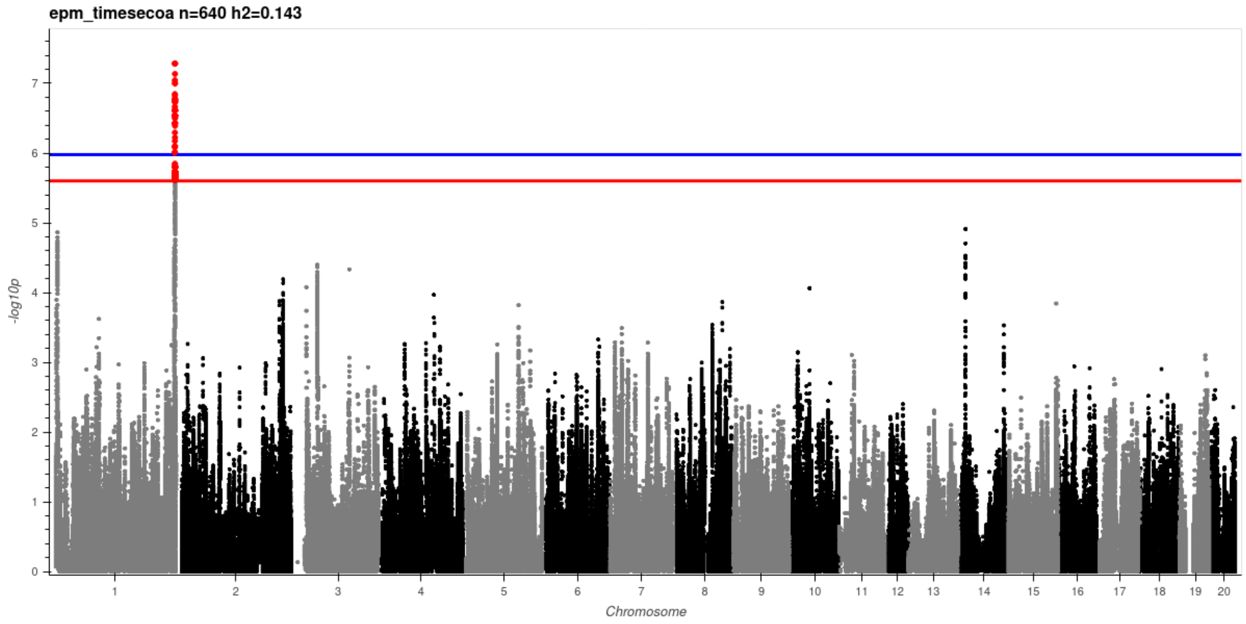


EPM Velocity

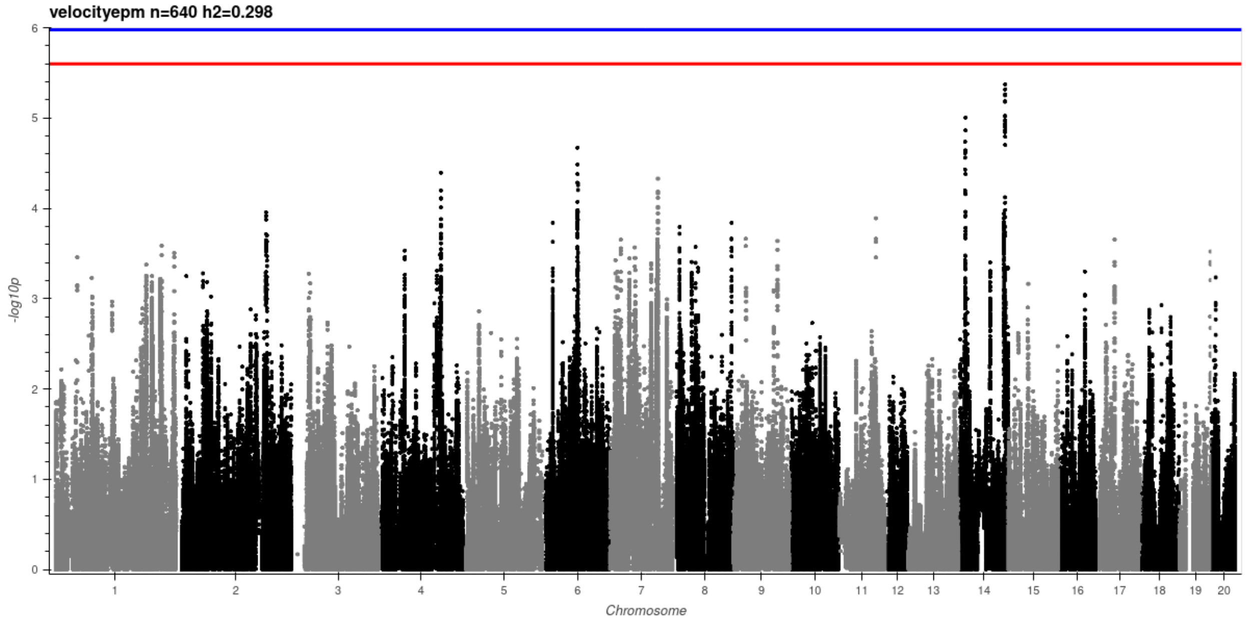

EPM Time Immobile

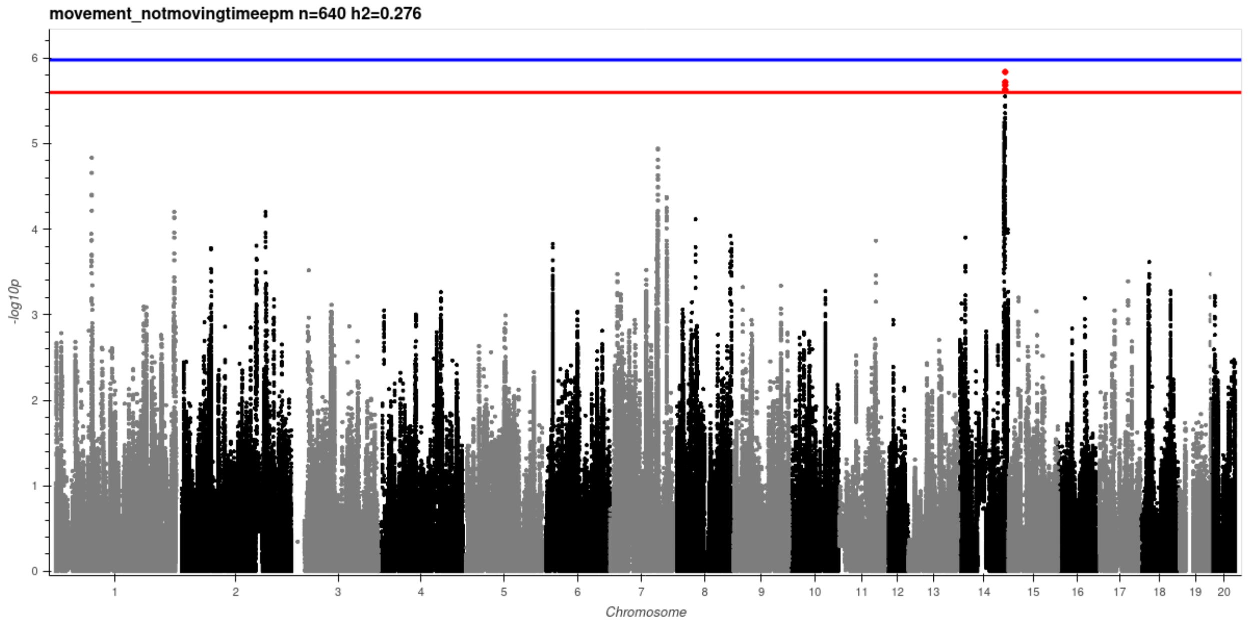


Tail Flick Latency

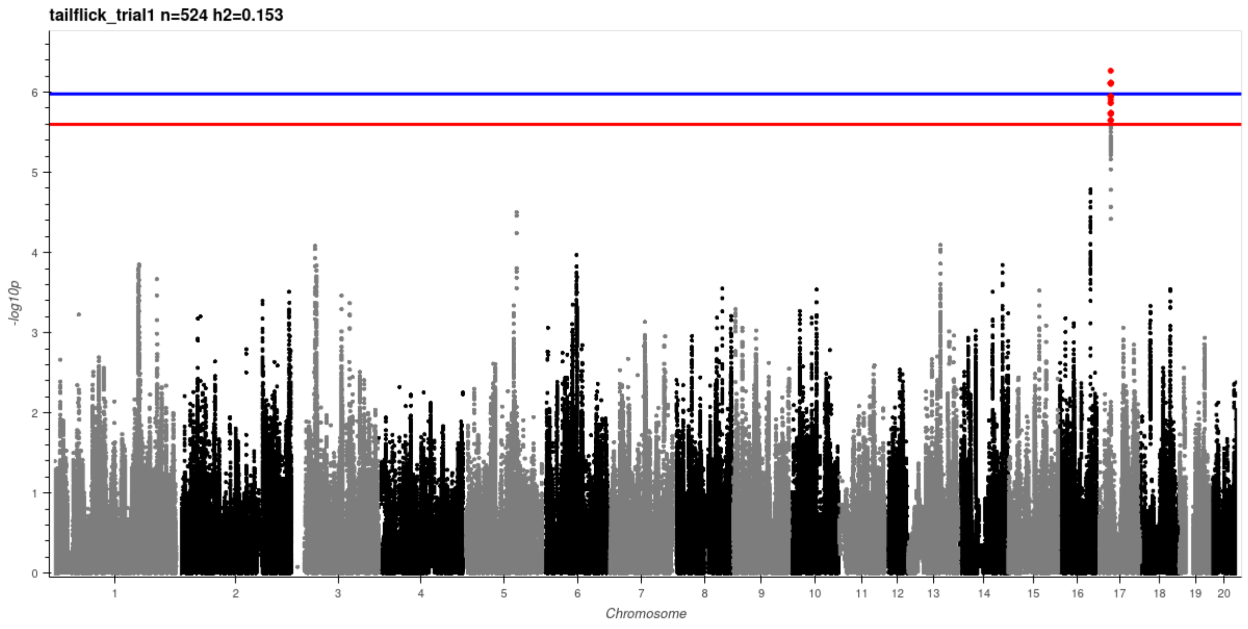

### Lateral Locomotor Score

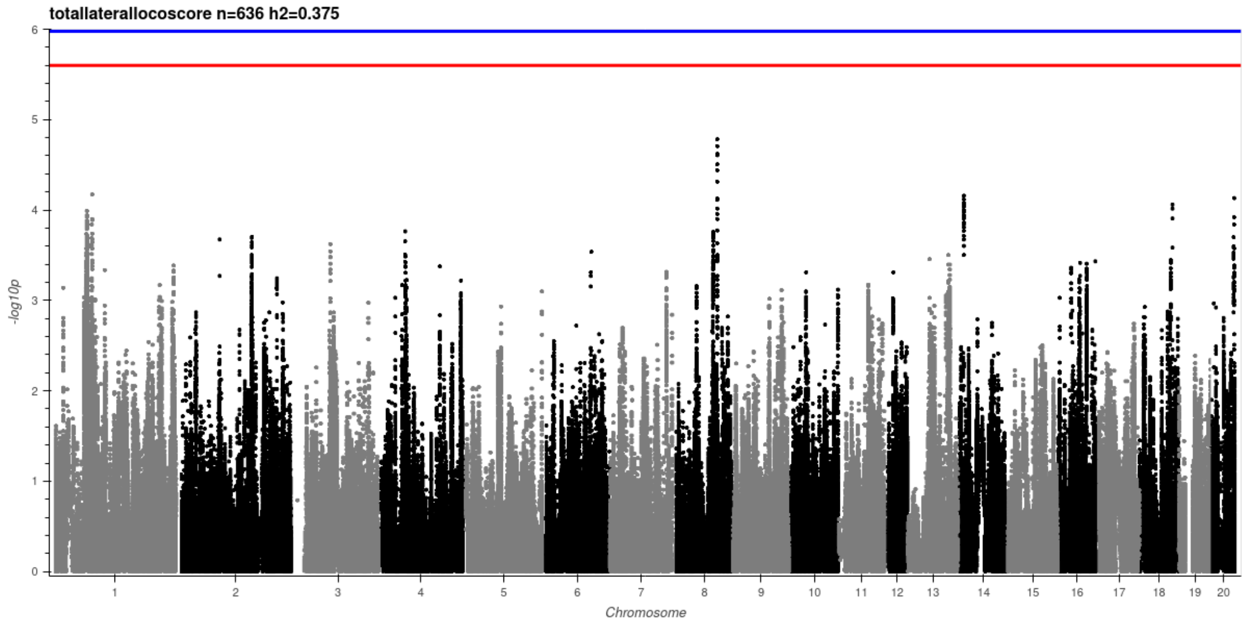


### Total Locomotor Score

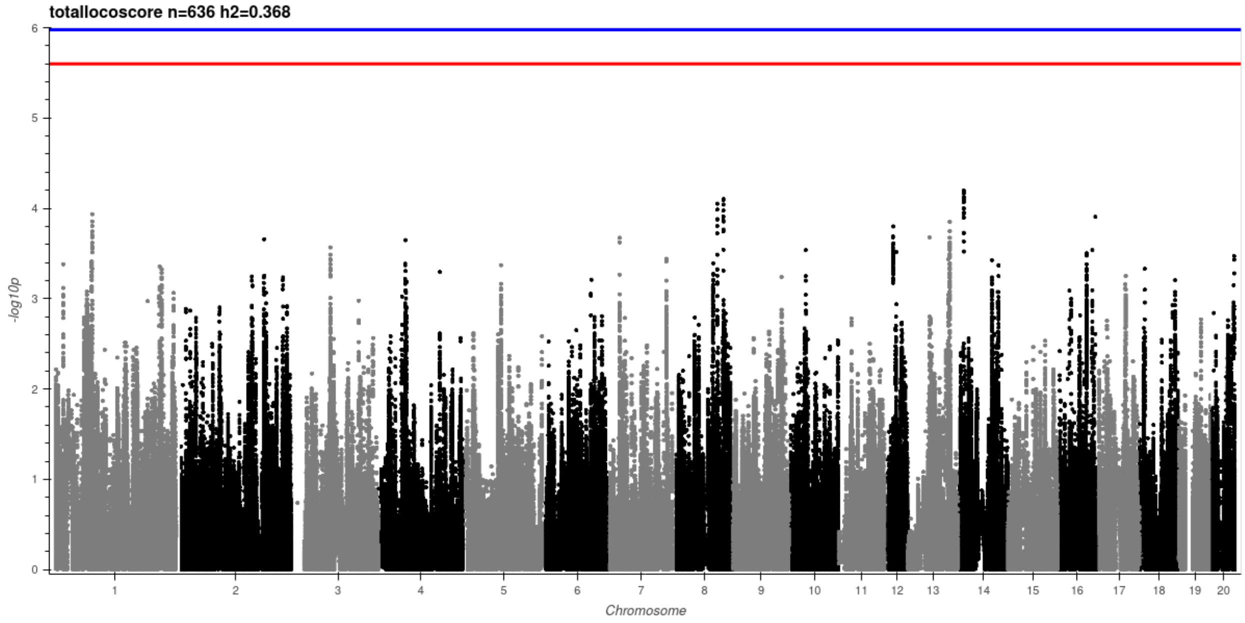

Rearing Locomotor Score

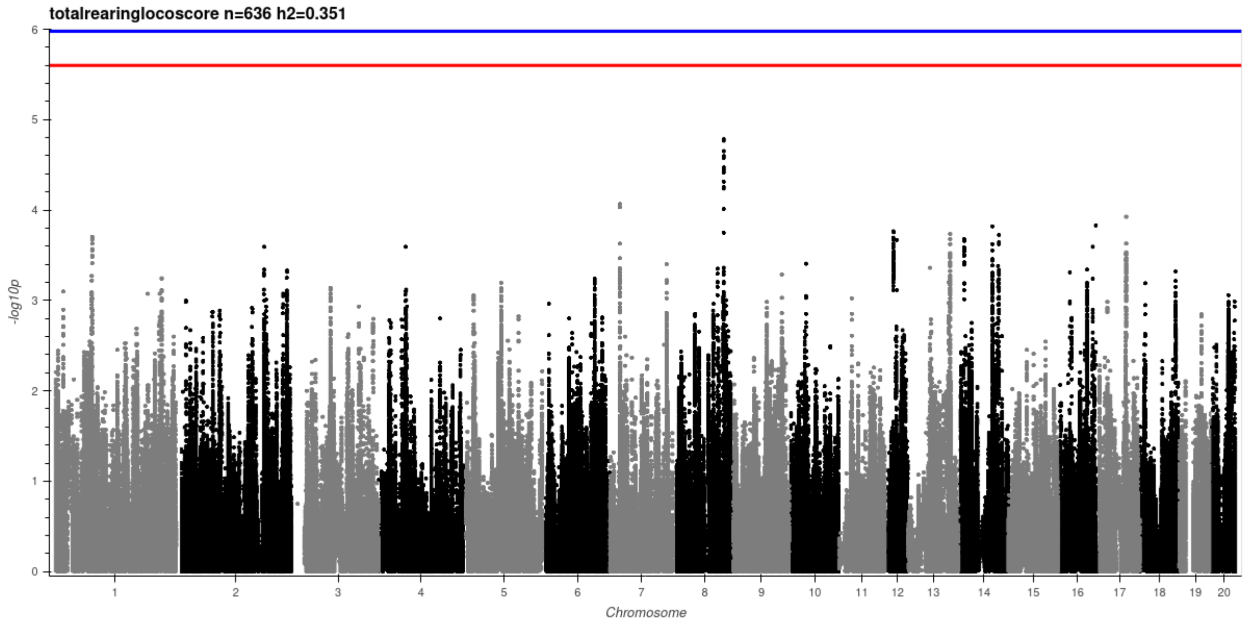

**Supplementary Figure S2.** Regional association plots for additional significant loci. A) EPM Open Arm Time Ratio at the chromosome 1 locus. B) EPM Open Arm Time at the same chromosome 1 locus. C) EPM Time Immobile at the chromosome 14 locus. Points are colored by LD r² to the lead SNP; axes show genomic position in Mb and −log10P. Horizontal lines mark the genome-wide and suggestive thresholds (−log10P = 5.98 and 5.60, respectively). The gene track displays annotated genes with transcription direction. Panels A and B map to the same chromosome 1 interval as the EPM Percent Time in Open Arms locus in the main figure.

A. EPM Open Arm Time Ratio at the Chromosome 1 Locus

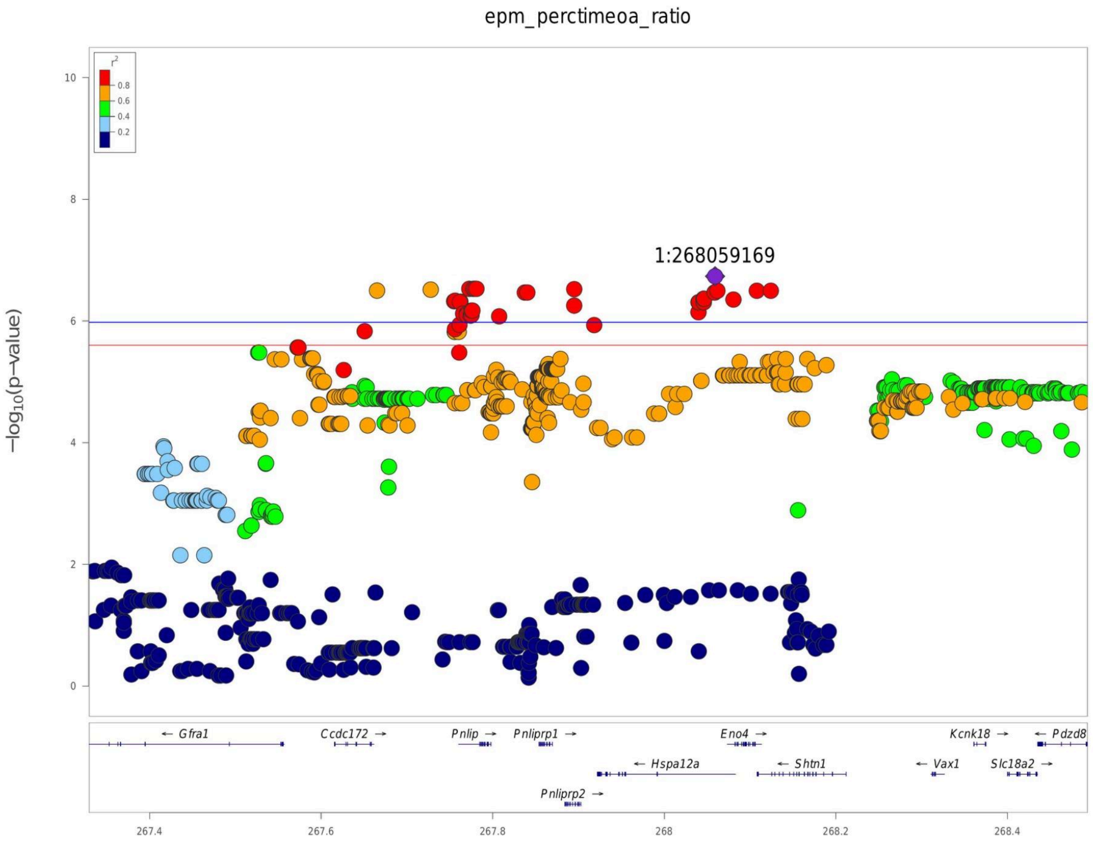

B. EPM Open Arm Time at the Chromosome 1 Locus

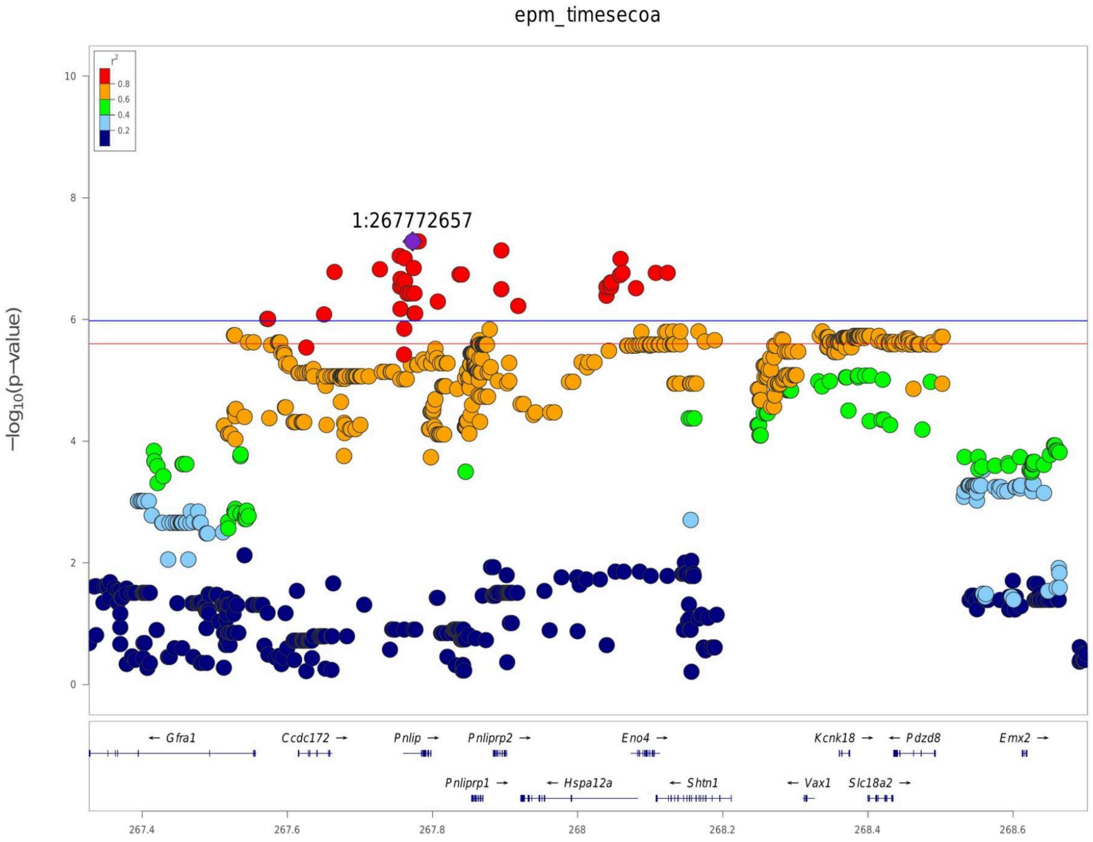

C. EPM Time Immobile at the Chromosome 14 Locus

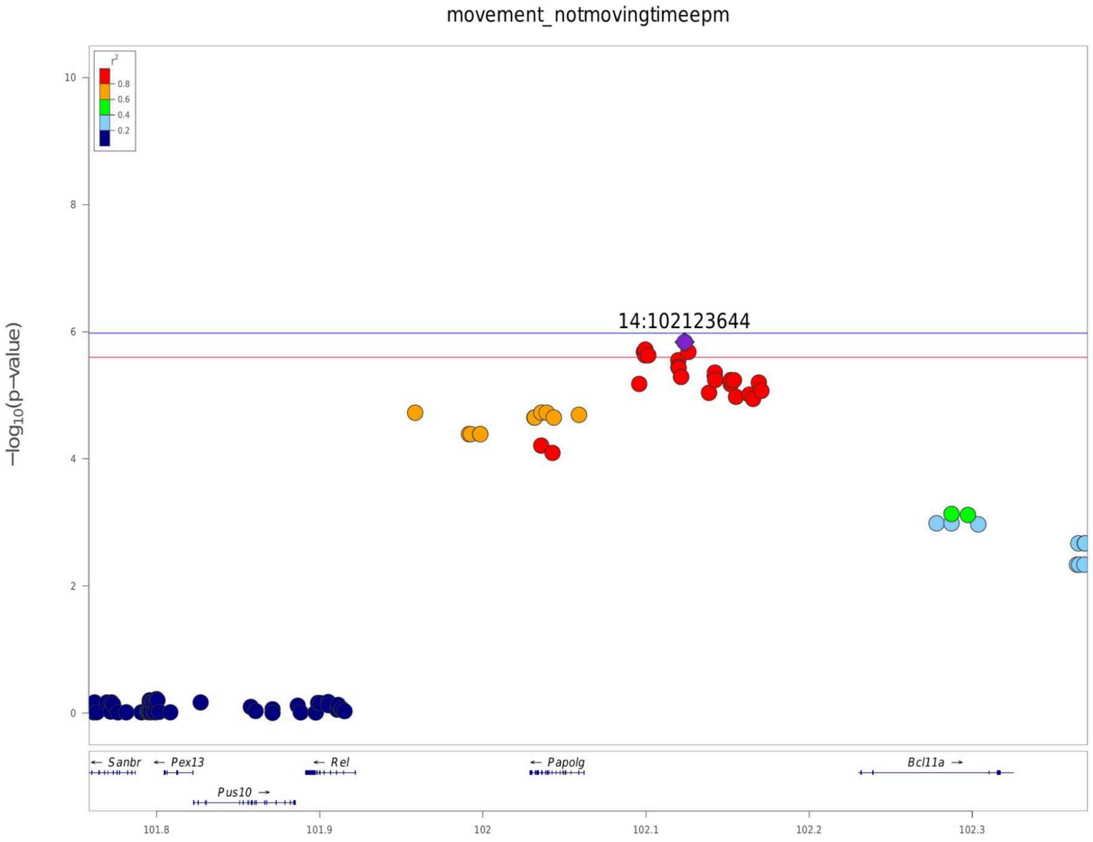
